# Implicating Gene and Cell Networks Responsible for Differential COVID-19 Host Responses via an Interactive Single Cell Web Portal

**DOI:** 10.1101/2021.06.07.447287

**Authors:** Kang Jin, Eric E. Bardes, Alexis Mitelpunkt, Jake Y. Wang, Surbhi Bhatnagar, Soma Sengupta, Daniel Pomeranz Krummel, Marc E. Rothenberg, Bruce J. Aronow

## Abstract

Numerous studies have provided single-cell transcriptome profiles of host responses to SARS-CoV-2 infection. Critically lacking however is a datamine that allows users to compare and explore cell profiles to gain insights and develop new hypotheses. To accomplish this, we harmonized datasets from COVID-19 and other control condition blood, bronchoalveolar lavage, and tissue samples, and derived a compendium of gene signature modules per cell type, subtype, clinical condition, and compartment. We demonstrate approaches to probe these via a new interactive web portal (http://toppcell.cchmc.org/ COVID-19). As examples, we develop three hypotheses: (1) a multicellular signaling cascade among alternatively differentiated monocyte-derived macrophages whose tasks include T cell recruitment and activation; (2) novel platelet subtypes with drastically modulated expression of genes responsible for adhesion, coagulation and thrombosis; and (3) a multilineage cell activator network able to drive extrafollicular B maturation via an ensemble of genes strongly associated with risk for developing post-viral autoimmunity.

## Introduction

COVID-19 clinical outcomes are variable. The poorer outcomes due to this infection are highly associated with immunological and inflammatory responses to SARS-Cov-2 infection (Shi et al., 2020; Tay et al., 2020) and many recent single cell expression profiling studies have characterized patterns of immunoinflammatory responses among individuals, mostly during acute infection phases. Different studies have revealed a spectrum of responses that range from lymphopenia (Cao, 2020; Wang et al., 2020), cytokine storms (Mehta et al., 2020; Pedersen and Ho, 2020), differential interferon responses (Blanco-Melo et al., 2020; Hadjadj et al., 2020) and emergency myelopoiesis (Schulte-Schrepping et al., 2020; Silvin et al., 2020). However, a variety of obstacles limit the ability of the research and medical communities to explore and compare these studies to pursue additional questions and gain additional insights that could improve our understanding of cell type specific responses to SARS-Cov-2 infection and their impact on clinical outcome.

Whereas many studies have focused on the peripheral blood mononuclear cells (PBMC) (Arunachalam et al., 2020; Guo et al., 2020; Lee et al., 2020; Schulte-Schrepping et al., 2020; Wilk et al., 2020a) due to ease of procurement, other studies have profiled airway locations via bronchoalveolar lavage (BAL) (Grant et al., 2020; Liao et al., 2020), nasopharyngeal swabs, and bronchial brushes (Chua et al., 2020). Additional sampling sites that could also be infected or affected have also been approached in autopsy-derived materials from the central nervous system (Heming et al., 2021; Yang et al., 2020), and other sites (Delorey et al., 2021). Moreover, as major COVID-19 consortiums working on the collection and integration of each of their individual studies and interpreting important features of these individual datasets as downloadable datasets or browsable versions, such as single cell portal (https://singlecell.broadinstitute.org/single_cell/covid19) and COVID-19 Cell Atlas (https://www.covid19cellatlas.org/), using these data beyond markers, cell types, and individual signatures is either not possible or not accomplishable across-datasets. Thus, a well-organized and systematic study of immune cells across tissues for in-depth biological explorations is an unmet need for a deeper understanding of the underlying basis of the breadth of COVID-19 host defense and pathobiology.

Here we harmonized and analyzed eight high quality publicly available single-cell RNA-seq datasets from COVID-19 and immunologically-related studies that in total covered more than 480,000 cells isolated from peripheral blood, bronchial alveolar lavage and lung parenchyma samples, and assembled an integrated COVID-19 atlas (https://toppcell.cchmc.org/). We established a framework for deriving, characterizing, and establishing reference gene expression signatures from these harmonized datasets using modular and hierarchical approaches based on signatures per class, subclass, and signaling/activation and clinical status per each sample group. Leveraging these gene expression signature modules, we demonstrate datamining approaches that allow for the identification of a series of fundamental disease processes: (1) an intercellular monocytic activation cascade capable of mediating the emergence of hyperinflammatory monocyte-derived alveolar macrophages in severe COVID-19 patients; (2) the generation of several alternatively differentiated platelet subtypes with dramatically different expression of sets of genes associated with critical platelet tasks capable of altering vascular and tissue responses to infectious agents; and (3) a multilineage and multi cell type cooperative signaling network with the potential to drive extrafollicular B maturation at a lesion site, but do so with high risk for the development of B cell-associated immunity. Additionally, immune hallmarks of COVID-19 patients were compared with other immune-mediated diseases using single-cell data from patients with influenza, sepsis, or multiple sclerosis. Consistent and varied compositional and gene patterns were identified across these implicating striking COVID-19 effects in some individuals.

## Results

### Creating the First COVID-19 Signature Atlas Using ToppCell Portal

To have a comprehensive coverage of cells, we collated single-cell data of COVID-19 patients from eight public datasets, which in total contains 231,800 PBMCs, 101,800 BAL cells and 146,361 lung parenchyma cells from donors: 43 healthy; 22 mild; 42 severe; and 2 convalescent patients (**Figure 1A, Table S1**).

**Fig. 1.**
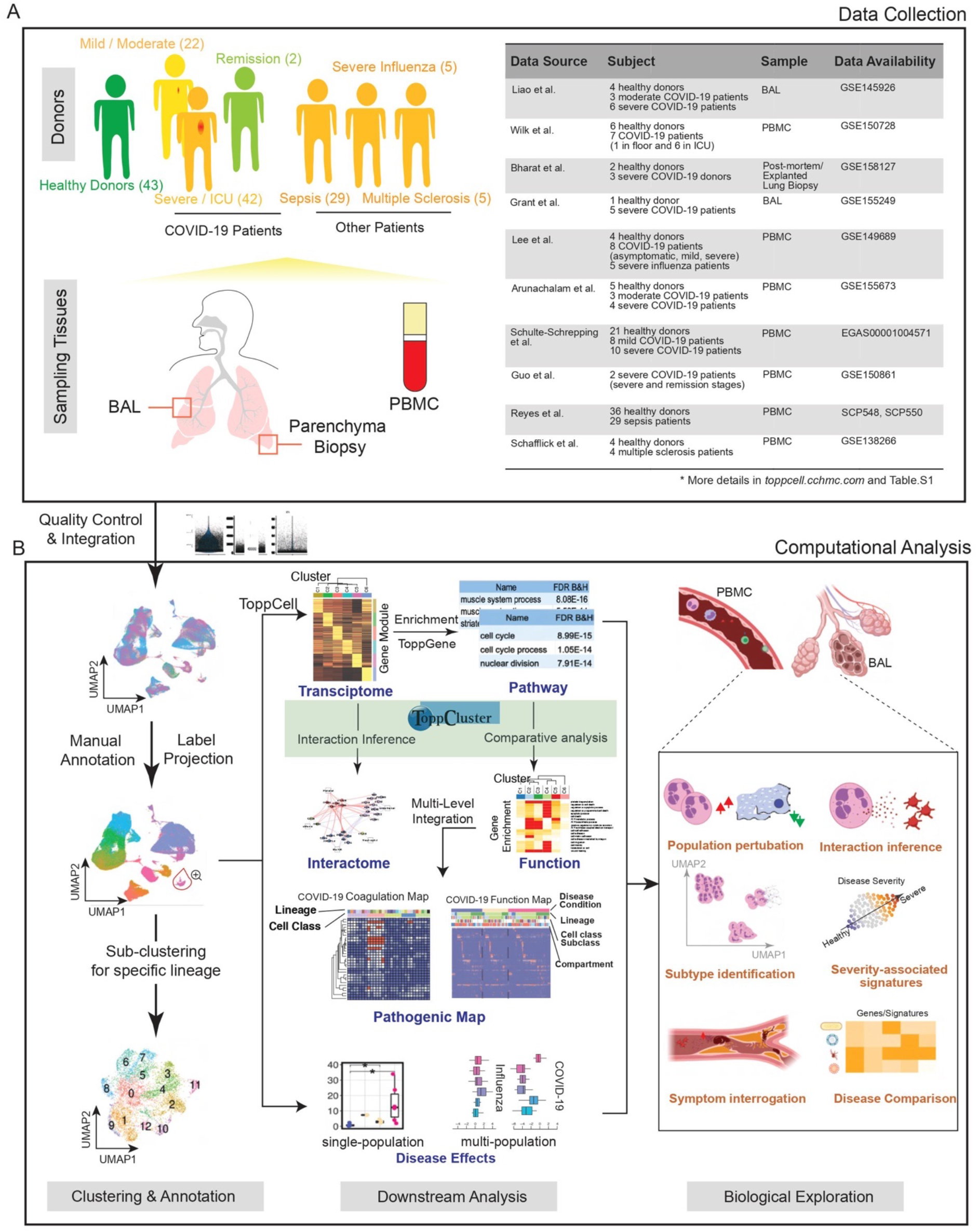
Creating a COVID-19 Signature Atlas. (**A**) Representative aggregation of multiple single-cell RNA-sequencing datasets from COVID-19 and related studies. The present study is derived from a total of 231,800 peripheral blood mononuclear cells (PBMCs), 101,800 bronchoalveolar lavage (BAL) cells and 146,361 lung parenchyma cells from 43 healthy; 22 mild, 42 severe, and 2 convalescent patients. Data was collated from eight public datasets (right). (**B**) Data analysis pipeline of the study using Topp-toolkit. It includes three phases: (1) clustering and annotation; (2) downstream analysis using Topp-toolkit; (3) biological exploration. Output includes the evaluation of abundance of cell populations, cell type (cluster) specific gene modules, functional associations of disease-associated cell classes and clusters, inference of cell-cell interactions, as well as comparative analysis across diseases, including influenza, sepsis and multiple sclerosis. Additional newer datasets not included in this manuscript are present and will continue to be added to ToppCell (http://toppcell.cchmc.org).

In order to assemble an integrated atlas of human cell responses to COVID-19, we sought to harmonize metadata encompassing clinical information, sampling compartments, and cell and gene expression module designations. Doing so provides a rich framework for detecting perturbations of cell repertoire and differentiative state adaptations. We first integrated single cell RNA-seq data in Seurat (Stuart et al., 2019) and annotated cell types using canonical markers (**Table S2**). Further annotations of B cell and T cell subtypes were completed using the reference-based labeling tool Azimuth (Hao et al., 2020). Sub-clustering was applied for some cell types, such as neutrophils and platelets, to interrogate finer resolutions of disease-specific sub-populations (**Figure 1B**). Using the ToppCell toolkit (https://toppcell.cchmc.org/), we created over 3,000 hierarchical gene modules of the most significant differentially expressed genes (DEGs) for all cell classes and sub-clusters across compartments and disease severity (**Table S1**). These modules were then used to infer cell-cell interactions as well as upregulated pathways, which were further combined for functional comparative analysis in a specific cell manner in ToppCluster (Kaimal et al., 2010) (**Figure 1B**), such as sub-clusters of platelets. Integration of ToppCluster output of cells from multiple compartments and disease conditions built pathogenic maps, highlighted by the coagulation map of COVID-19 (**Figure S12**). In addition, perturbation of cell abundance was evaluated either in one cell population, or in multiple populations across diseases. Taken together, we investigated cell abundance changes, severity-associated signatures, mechanisms of COVID-19 specific symptoms and unique features of COVID-19 as an immune-mediated disease (**Figure 1B**).

### Dynamic Changes and Balance of COVID-19 Immune Repository in Blood and Lung

After the aforementioned cell annotation procedure, we identified 28 and 24 distinct cell types in PBMC and BAL respectively (**Figure 2A, 2C; Table S2**). Shifts of Uniform Manifold Approximation and Projection (UMAP) of cell type distributions were observed in both compartments of mild and severe patients (**Figure 2A, 2C, S1A and 3A**). In PBMC, conventional dendritic cells (cDC), plasmacytoid dendritic cells (pDC) and non-classical monocytes displayed a prominent reduction in severe patients (**Figure 2B, S1C**), consistent with prior reports (He et al., 2020; Laing et al., 2020; Wilk et al., 2020a). In contrast, severe patients demonstrated dramatic expansion of neutrophils, especially immature stages (**Figure S1C, S2**). Integration with evoked pathways in the following analysis implicated that neutrophil expansion was likely the consequence of emergency myelopoiesis (Wilk et al., 2020b). Additionally, a general down-regulation of T cell and NK cell was observed, consistent with lymphopenia reported in clinical practices (Pedersen and Ho, 2020; Terpos et al., 2020) (**Figure S1C, S2**). However, the trend of T cell subtypes varies across studies and individuals, apart from proliferative T cells which have a dramatic increase in mild and severe patients (**Figure S2**). Notably, plasmablasts substantially increased in COVID-19 patients, and especially so in severe patients, suggesting upregulated antibody production (De Biasi et al., 2020) (**Figure 2B and S1C**). Expansion of platelets is another significant change observed in severe patients, possibly leading to immunothrombosis in the lung, which could be closely associated with the severity of the disease (Middleton et al., 2020; Nicolai et al., 2020) (**Figure 2B and S1C**).

**Fig. 2.**
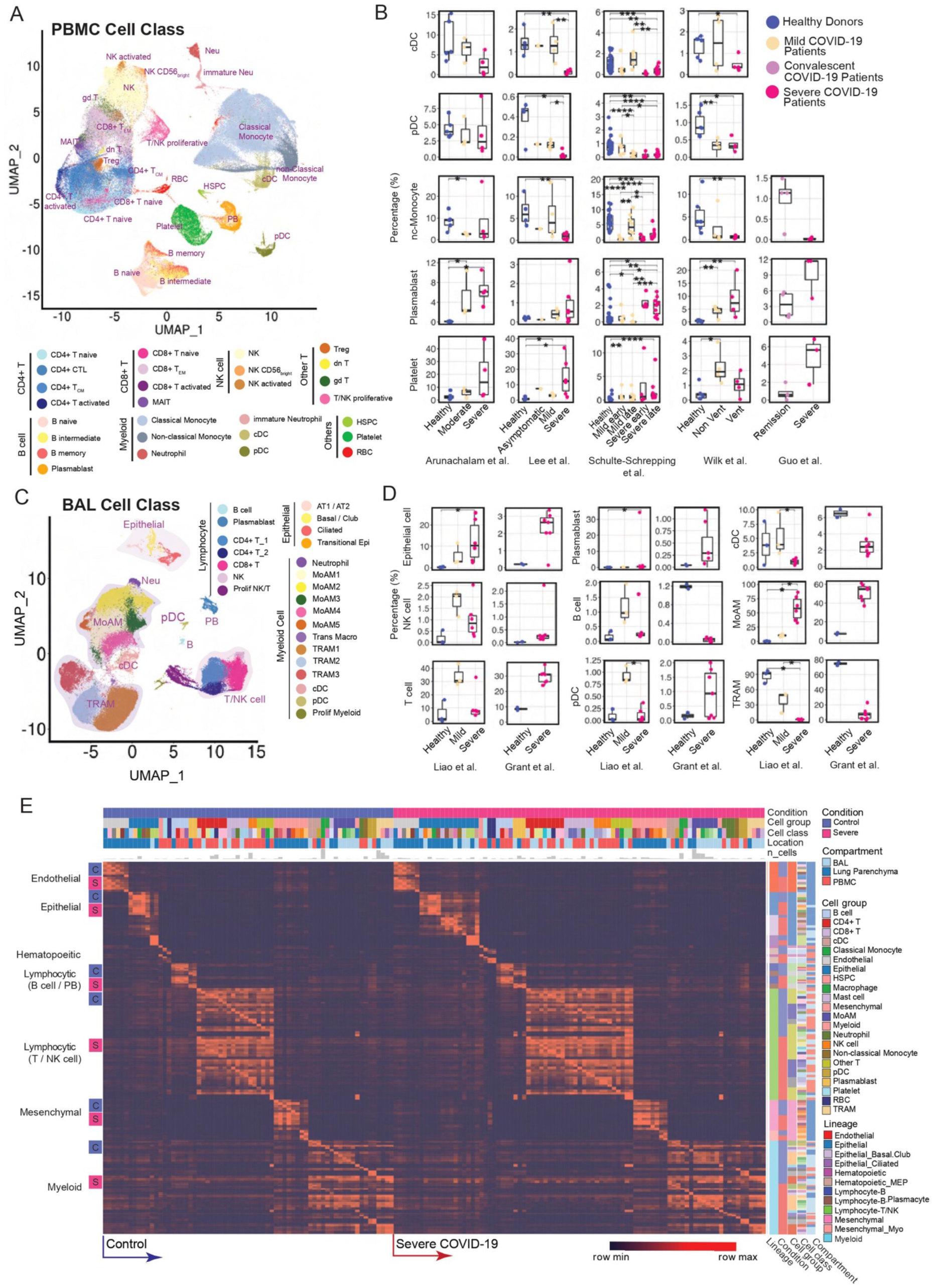
Modularized representation of cell type specific gene signatures and dynamic changes of cell abundance. (**A**) Uniform Manifold Approximation and Projection (UMAP) of 28 distinct cell types identified in the integrated peripheral blood mononuclear cell (PBMC) data. (**B**) Comparative analysis of cell abundance effects of COVID-19. Reproducible multi-study data present high impact effects on 5 cell types in PBMC. Percentages of selected cell types in each sample are shown (where Vent: Ventilated patients; Non Vent: Non-ventilated patients). Significance between two conditions was measured by the Mann-Whitney rank sum test (Wilcoxon, paired=False), which was also used in following significance tests of cell abundance changes in this study. *: p <= 0.05; **: p <= 0.01; ***: p <= 0.001; ****: p <= 0.0001. (**C**) UMAP of 24 distinct cell types identified in the integrated BAL data. (**D**) Dynamic changes of cell abundances for cell types in two bronchoalveolar lavage (BAL) single-cell datasets. (**E**) ToppCell allows for gene signatures to be hierarchically organized by lineage, cell type, subtype, and disease condition. The global heatmap shows gene modules with top 50 upregulated genes (student t test) for each cell type in a specific disease condition and compartment. Gene modules from control donors and severe COVID-19 patients were included in the figure.

In samples obtained from patients’ lungs, we observed the depletion of FABP4^high^ tissue-resident alveolar macrophages (TRAM) and dramatic expansion of FCN1^high^ monocyte-derived alveolar macrophages (MoAM) in severe patients (**Figure 2C, 2D and S3D**). Mild patients exhibited a moderate reduction of tissue-resident macrophages, but no evidence of aggregation of monocyte-derived macrophages (**Figure 2C, 2D, S3A and S3D**). Dynamic changes of these two subtypes suggest increased tissue chemoattraction (Merad and Martin, 2020) and potential damage of patients’ lungs (McGonagle et al., 2020). In addition, neutrophils were only identified in severe patients in the integrated BAL data (**Figure 2C and S3A**), which might be related with neutrophil extracellular traps (NETs) in the lung (Barnes et al., 2020). However, more samples are required to draw a solid conclusion. We also noted conventional dendritic cells decreased in the severe patients, which is consistent with the trend of the counterpart in PBMC data. Opposite to the change in PBMC, an expansion of plasmacytoid dendritic cells is observed in both mild and severe patients (**Figure 2D**). Other cell types, including T cell and NK cell in the BAL, also have converse changes of their counterparts in PBMC, which could be attracted by lung macrophages or epithelial cells after infection or damages (Chua et al., 2020) (**Figure 2D and Figure S3D**). These changes were consistently observed in lung parenchyma samples from severe COVID-19 patients (**Figure S4**). With cells well-annotated in the integrated COVID-19 atlas, we drew a global heatmap for cells in both blood and lung using ToppCell gene modules (top 50 DEG in each module) of all identified cell classes. While there was conservation of gene patterns involved in healthy donors and severe COVID-19 patients, there were substantial differences most notably in myeloid cells (**Figure 2E**). Such hierarchically ordered ToppCell gene modules were broadly used in visualization, large-scale comparisons and fine-resolution investigations in the following analyses.

### Myeloid Cell Atlas: Functionally Distinct Neutrophils at Different Levels of Maturation and Derailed Macrophages in the Lung

Dysregulated myeloid cells have been reported as an important marker of severe COVID-19 patients (Schulte-Schrepping et al., 2020; Silvin et al., 2020). In order to gain a deeper and comprehensive understanding of these cells, we applied the sub-clustering strategy on the integrated data of key cell types, such as neutrophils and macrophages, and then generated gene modules for comparative functional analysis and interactome inference. We successfully identified 5 neutrophil sub-clusters after the integration of PBMC and BAL data, including 3 FCGR3B+ mature sub-clusters and 2 FCGR3B-immature sub-clusters (**Figure 3A and S5B; Table S3**). They’re mainly from severe patients and their gene modules were generated and subjected to comparative functional enrichment using ToppCell and ToppCluster (**Figure 3C, 3D and S5A**). We identified proliferative neutrophils (referred to as pro-neutrophils and Neu4) and MMP8^high^ precursor immature neutrophils (referred to as pre-neutrophils and Neu2) (**Figure 3A and S5B**) consistent with prior studies (Schulte-Schrepping et al., 2020). While immune response genes and pathways were barely activated in the immature neutrophils, they displayed upregulation of granule formation pathways and NETosis-associated proteins, including ELANE, DEFA4 and MPO, especially in Neu4 (Schulte-Schrepping et al., 2020; Wilk et al., 2020b) (**Figure 3C, S5B and S5C**). Upregulated myeloid leukocyte mediated immunity in Neu2 suggests involvement of this cell type in anti-viral function (**Figure S5D**). Yet, the absence of cytokine and interferon response pathways suggests the lack of mature immune responses (**Figure 3D**). Notably, compared to mature neutrophils (Neu0 and Neu1) in the blood, the extravasated hyperinflammatory sub-cluster (Neu3) from BAL of severe patients shows extraordinarily high expression of interferon-stimulated genes, as well as prominent upregulation of productions and responses to cytokines and interferons (**Figure 3C, 3D, S5B and S5D**).

**Fig. 3.**
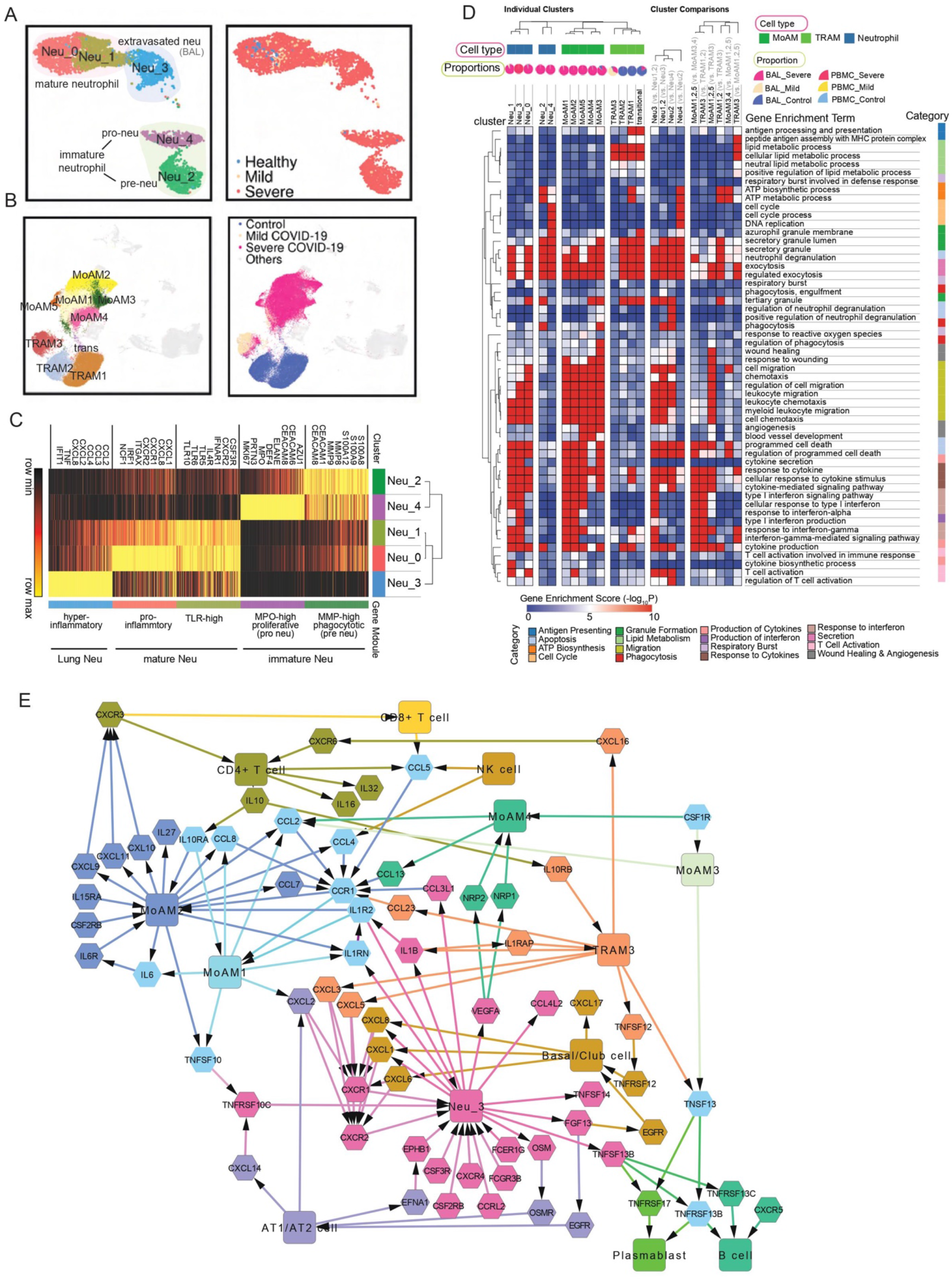
Functional analysis of compartment-specific immature and subtype-differentiated neutrophils and monocytic macrophages in COVID-19 patients. (**A**) Five sub-clusters and three cell groups were identified after the integration of neutrophils in peripheral blood mononuclear cells (PBMC) and bronchoalveolar lavage (BAL) (Left). The distribution of compartments is shown on the right. (**B**) Sub-clusters (Left) and COVID-19 conditions (Right) of monocyte-derived macrophages and tissue-resident macrophages were identified after integration of BAL datasets. (**C**) Heatmap of gene modules from ToppCell with top 200 upregulated genes for each neutrophil sub-cluster. Important neutrophil-associated genes and inferred roles of sub-clusters were shown on two sides. (**D**) Heatmap of associations between subclusters of neutrophils and macrophages and myeloid-cell-associated pathways (Gene Ontology). Gene modules with 200 upregulated genes for sub-clusters were used for enrichment in ToppCluster. Additionally, enrichment of top 200 differentially expressed genes (DEGs) for comparisons in fig. S5D and fig. S6B were appended on the right. Gene enrichment scores, defined as -log_10_(adjusted p-value), were calculated as the strength of associations. Pie charts showed the proportions of COVID-19 conditions in each cluster. (**E**) Gene interaction network in the BAL of severe patients. Highly expressed ligands and receptors of each cell type were drawn based on fig. S8. Interaction was inferred using both CellChat database and embedded cell interaction database in ToppCell.

MoAM and TRAM were two main macrophage types in the BAL (**Figure 2C**); both are known to have distinct roles in immune responses in the lung (Liao et al., 2020). As described above, five sub-clusters among the expanded COVID-19 patient-specific MoAM (**Figure 3B; Table S3**) were found, where the loss of HLA class II genes and elevation of interferon-stimulated genes (ISGs) were consistently observed (**Figure 3F and S6A**). Relative to MoAM3,4, MoAM1,2,5 displayed an upregulation of interferon responses and cytokine production (**Figure 3D and S6B; Table S3**), indicating their pro-inflammatory characteristics. Notably, MoAM5 shows dramatic upregulation of IL-6 secretion and cytokine receptor binding activities (**Figure S7A-D**). However, cells in this sub-cluster were mainly from one severe patient (**Figure S3C**). We still need more data to fully understand such dramatic upregulation of IL-6 secretion in some severe patients. Similar to MoAM, we also identified two distinct groups of TRAM in BAL (**Figure 3B and S6B**), including quiescent TRAM (TRAM1 and TRAM2) and activated TRAM (TRAM3). The quiescent group was mainly from healthy donors with enriched pathways of ATP metabolism (**Figure 3D**), while the activated group from mild and severe patients displays upregulation of ISGs and cytokine signaling pathways (**Figure S6B; Table S3**). However, the magnitude of activation and inflammatory responses in TRAM3 is smaller than MoAM1,2,5. Not surprisingly, stronger antigen processing and presentation activities were observed in TRAM3 relative to MoAM1,2,5 (**Figure 3D and S6B; Table S3**). Collectively, we concluded that tissue-resident macrophages were greatly depleted in severe patients as the front-line innate immune responders in the lung. Pro-inflammatory monocyte-derived macrophages infiltrate into the lung, leading to the cytokine storm and damage of the lung. Large amounts of infiltration of MoAM were not observed in mild COVID-19 patients, probably due to the controlled infection, which could explain milder lung damages in those patients.

To develop an understanding of the interaction network in the lung microenvironment of severe COVID-19 patients, we focused on signaling ligands, receptors and pathways using ToppCell and CellChat (**Figure 3E, S8A, S8B**). Notably, basal cells, MoAMs, neutrophils and T cells all contributed to the cytokine, chemokine and interleukin signaling networks. Strikingly, severe patient specific MoAM2 shows the broadest upregulation of signaling ligands, including CCL2, CCL3, CCL7, CCL8, CXCL9, CXCL9, CXCL10, CXCL11, IL6, IL15 and IL27, suggesting its role as a signaling network hub that is distinct from the other major signaling ligand-expressing cells of BAL such as epithelial and other myeloid cell types such as TRAM3 and proliferating myeloid cells (**Figure S8A**). Among the MoAM2 top signaling molecules, attractants CXCL8, CXCL9 and CXCL10 are known to target CXCR3 on T cells, suggesting their role is to stimulate migration of T cells to the epithelial interface and into BAL fluid (**Figure 3E**) (Liao et al., 2020). In addition, many of MoAM2’s ligands have the potential to cause autocrine signaling activation via IL6-IL6R, IL1RN-ILR2, CCL7-CCR1, CCL2-CCR1 and CCL4-CCR1, indicating its active roles in self-stimulation and development, which further amplify the attraction and migration of T cells and other immune cells. Notably, CCR1 was also expressed in activated TRAM3, but with a lower level. Although IL6 expression level is relatively low compared to other ligands in BAL data, substantial expression of IL6R was observed in MoAMs. The CCL and CXCL signaling pathways of neutrophils are less strong than MoAMs (**Figure S8B**), but they displayed high expression levels of CXCR1 and CXCR2, which binds with a large number of the chemokines from MoAM and epithelial cells (**Figure 3E**). In addition, neutrophils exhibit an extraordinarily high level of IL1B, which could potentially in turn activate macrophages (**Figure S8A and S8B**). TRAM3 also displayed a unique pattern of signaling molecules, with a substantial level of CCL23 which could potentially attract MoAM by the interaction with CCR1. Secretion of CXCL3 and CXCL5 in TRAM3 towards CXCR2 could be a potential chemoattraction pathway for neutrophils. In turn, neutrophils could activate TRAM3 by secreting IL1B, which binds with IL1RAP. Additionally, CD4+ T cells could also activate TRAM3 by IL10-IL10RB interaction (**Figure 3E, S8A and S8B**).

In addition to neutrophils and macrophages, the upregulation of ISGs was observed in classical monocytes of both mild and severe patients (cMono3, cMono4), while the reduction of the MHC class II cell surface receptor HLA-DR genes was only observed in severe patients (cMono4) (**Figure S9**). In cDCs, polarization of interleukin secretion was observed in mild-patient and severe-patient specific clusters (**Figure S10F**). Collectively, dynamic changes of marker genes, transcriptional profiles, signaling molecules and biological activities reveal the heterogeneity of myeloid cell sub-clusters across disease severity (**Figure S11C**). Pro-inflammatory gene expression was found in all major myeloid cell types, including cMono4, DC1, DC9 in PBMC and Neu3, DC10, MoAM1, MoAM2, MoAM5 in BAL of COVID-19 patients. The reduction of MHC class II (HLA-II) genes is a common feature of classical monocytes and macrophages in COVID-19 patients and implies impaired capacity to activate T cell adaptive immunity.

### COVID-19 Coagulation and Immunothrombosis Map

Individuals severely affected during acute phase COVID-19 infection, and in particular those with significantly elevated risk of death, frequently demonstrate striking dysregulation of coagulation and thrombosis characterized by hypercoagulability and microvascular thromboses (endothelial aggregations of platelets and fibrin) and highly elevated D-dimer levels. Yet, COVID-19 does not lead to wide scale consumption of fibrinogen and clotting factors (Iba et al., 2020a; Levi et al., 2020; Middleton et al., 2020; Nicolai et al., 2020; Rapkiewicz et al., 2020). At present, we lack a molecular or cellular explanation of the underlying basis of this pathobiology (Aid et al., 2020; Middleton et al., 2020). To evaluate candidate effectors of this pathobiology, we used a list of genes associated with abnormal thrombosis from mouse and human gene mutation phenotypes and identified parenchymal lung sample endothelial cells and platelets in PBMC as cell types highly enriched with respect to genes responsible for the regulation of hemostasis (**Figure S12**). Because platelet counts were greatly elevated in severe versus mild individuals, we further examined platelet gene expression signatures and cell type differentiation and identified six distinct platelet sub-clusters shared across all datasets after data integration (**Figure 4A and 4B**). Severe-patient-specific PLT0 is an interesting sub-cluster with elevated integrin genes, including ITGA2B, ITGB1, ITGB3, ITGB5, as well as thrombosis-related genes, such as SELP, HPSE, ANO6 and PF4V1. Antibodies against the latter are associated with thrombosis including adverse reactions to recent COVID-19 vaccine ChAdOx1 nCoV-19 (Schultz et al., 2021). In addition, upregulated pathways of hemostasis, wound healing and blood coagulation were also observed in PLT0 (**Figure S13A; Table S4**). Importantly, PLT2 is an inflammatory sub-cluster with an upregulation of ISGs and interferon signaling pathways, while PLT4 is highlighted by upregulated post-transcriptional RNA splicing activities (**Figure S13A and S13C**).

**Fig. 4.**
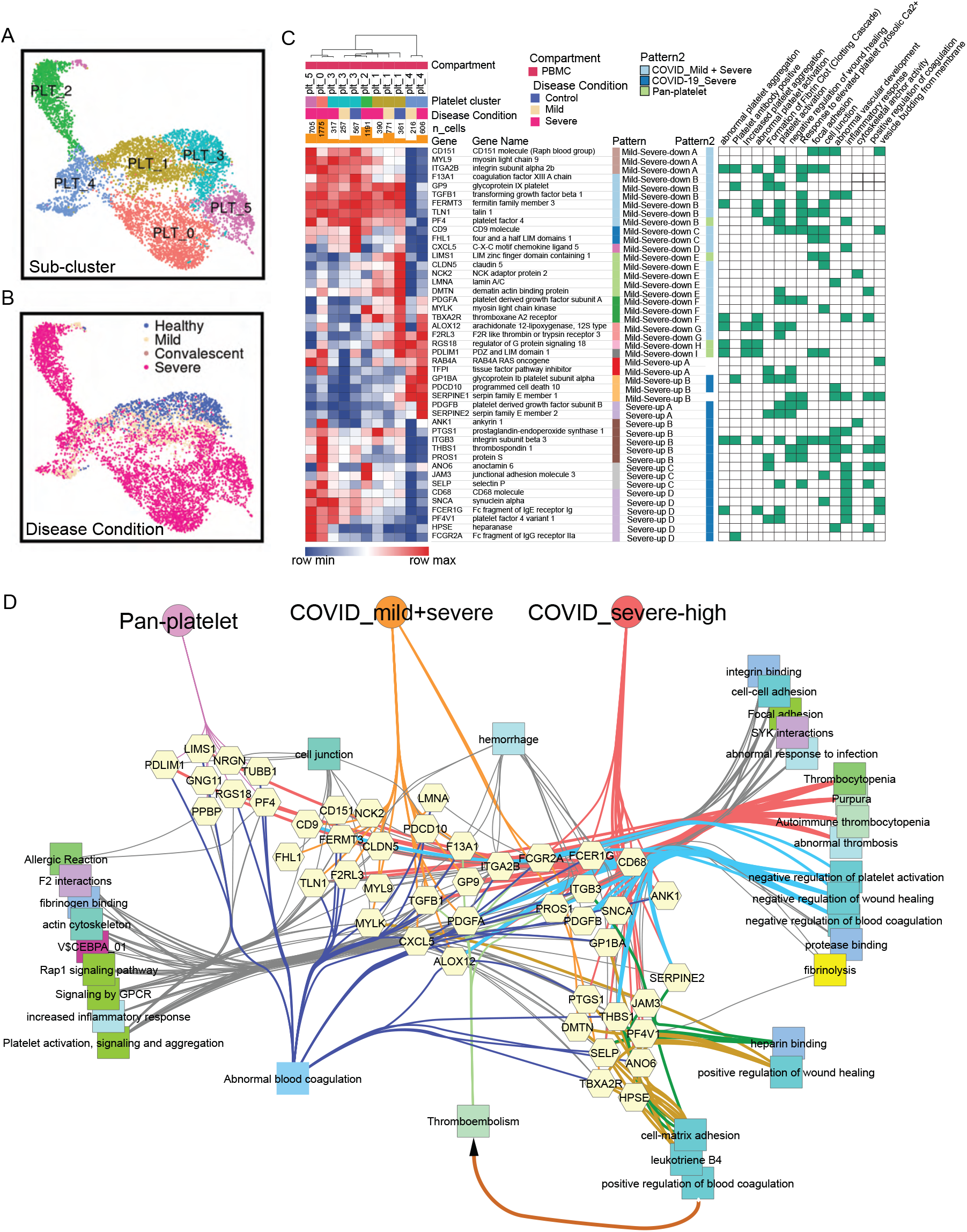
COVID-19 driven reprogramming of platelets leads to drastically altered expression of genes associated with platelet adhesion, activation, coagulation and thrombosis. (**A-B**) Uniform Manifold Approximation and Projections (UMAPs) show distributions of sub-clusters (A) and COVID-19 conditions (B) of platelets after the integration of PBMC datasets. (**C**) Severity-associated coagulation genes were selected and shown on the heatmap, with disease and sub-cluster specific gene patterns identified and labeled. Their functional associations with coagulation pathways were retrieved from ToppGene and shown on the right. (**D**) Functional and phenotypical associations of coagulation-association genes in each gene pattern from (B). Associations were retrieved from ToppGene enrichment. Fibrinolysis is highlighted.

Severity-associated gene patterns were also identified by selecting coagulation-associated genes modules (**Figure 4C; Table S4**), indicating distinct coagulation activities across platelets. Apart from pan-platelet genes, we found dramatic upregulation of genes involved in platelet activation, fibrinogen binding and blood coagulation in platelets of severe COVID-19, including procoagulant heparanase (HPSE) (Osterholm et al., 2013), Anoctamin-6 (ANO6) (Swieringa et al., 2018), and selectin P (SELP) (Sparkenbaugh and Pawlinski, 2013) (**Figure 4C and 4D**). Heparanase is an endoglycosidase that cleaves heparan sulfate constituents, a major component of anti-coagulation glycocalyx on the surface of vascular endothelium (Edovitsky et al., 2006; Iba et al., 2020b). Upregulated heparanase was related to upregulation of cell-matrix adhesion and coagulation (**Figure 4D**). Thrombotic vascular damages could be caused by the degradation function of heparinase enriched in platelets of severe patients. Elevation of ANO6 is known to trigger phospholipid scrambling in platelets, resulting in phosphatidylserine exposure which is essential for activation of the clotting system (Heemskerk et al., 2013). In addition, other upregulated genes involved in coagulation-associated activities were also observed, including wound healing, fibrinolysis, platelet aggregation and activation (**Figure 4D**), which likely collectively contribute to the clotting issue of severe COVID-19 patients.

### Emergence of Developing Plasmablasts and B Cell Association with Autoimmunity

Autoimmune disorders in COVID-19 patients such as immune thrombocytopaenic purpura (ITP) is now recognized as a known disease complication (Ehrenfeld et al., 2020; Fujii et al., 2020; Gruber et al., 2020; Rodríguez et al., 2020; Zhou et al., 2020). However, little is known about the molecular and cellular mechanism behind it. To examine this further, we integrated B cells and plasmablasts from both PBMC and BAL and conducted systematic analysis (**Figure 5A, 5B, S14A and S14B**). Several COVID-19 specific sub-clusters were identified in B cells, such as ISG^high^ activated B cells (cluster 7) (**Figure 5A**). Importantly, activated B cells showed dramatic upregulation of interferon signaling pathways and cytokine productions (**Figure S15A**), indicating its anti-virus characteristics. Notably, plasmablasts were mainly observed in severe COVID-19 patients, where a group of proliferative cells was identified and labeled as developing plasmablasts (**Figure 5B**). In contrast, non-dividing plasmablasts displayed upregulation of immunoglobulin genes (IGHA1, IGHA2, IGKC), B cell markers (CD79A) (Mason et al., 1995), interleukin receptors (IL2RG) and type II HLA complex (HLA-DOB) (**Figure 5C; Table S5**). In addition, non-dividing plasmablasts showed unique isotypes of immunoglobulin (Ig) in sub-regions of UMAP, whereas developing plasmablasts displayed obscure Ig types (**Figure S14E and S14F**). Antibody production activities were upregulated in non-dividing plasmablasts based on gene enrichment analysis (**Figure S15A; Table S5**). Collectively, we inferred that non-dividing plasmablasts had definite immunoglobulin isotypes and were actively involved in immune responses towards COVID infection, while developing plasmablasts were less mature but highly proliferative to replenish the repertoire of plasma cells.

**Fig. 5.**
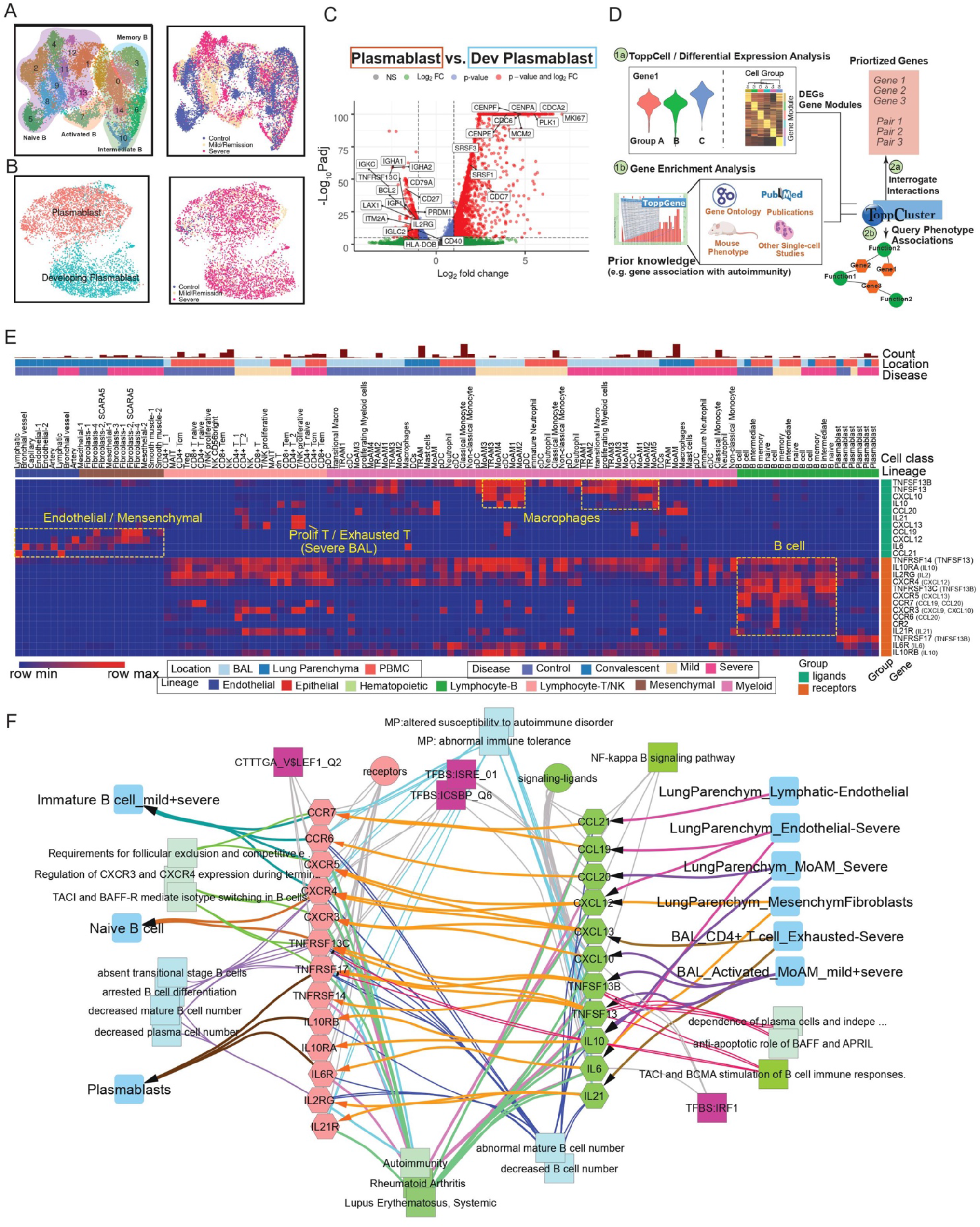
Implicating a multi-lineage cell network capable of driving extrafollicular B cell maturation and the emergence of humoral autoimmunity in COVID-19 patients. (**A**) Uniform Manifold Approximation and Projections (UMAPs) of sub-clusters (Left) and COVID-19 conditions (Right) of B cells after integration of peripheral blood mononuclear cells (PBMC) and bronchoalveolar lavage (BAL) datasets. (**B**) UMAPs of subtypes (Left) and COVID-19 conditions (Right) of plasmablasts after integration of PBMC and BAL datasets. (**C**) Volcano plot depicts differentially expressed genes between plasmablasts and developing plasmablasts. Student t-tests were applied and p values were adjusted by the Benjamini-Hochberg procedure. (**D**) Workflow of discovering and prioritizing candidate genes related to a disease-specific phenotype with limited understanding. (**E**) The heatmap shows the normalized expression levels of candidate ligands and receptors for COVID-19 autoimmunity in multiple compartments in healthy donors and COVID-19 patients. Binding ligands of receptor genes were shown in parentheses on the right. Hot spots of expression are highlighted. (**F**) Network analysis of autoimmunity-associated gene expression by COVID-19 cell types. Prior knowledge associated gene associations include GWAS, OMIM, mouse knockout phenotype, and additional recent manuscripts were selected from ToppGene enrichment results of differentially expressed ligands and receptors and shown on the network. Orange arrows present the interaction directions from ligands (green) to receptors (pink) on B cells. Annotations for these genes, including single-cell co-expression (blue), mouse phenotype (light blue), transcription factor binding site (purple) and signaling pathways (green) are shown.

Since there are few clues of gene associations of autoimmunity in COVID-19, we brought up a hypothesis-driven, prior knowledge-based approach to discover and prioritize genes for the specific phenotype (**Figure 5D**). First, gene modules of B cells and other cells in severe patients were collected and subjected to ToppGene for enrichment analysis. Then we queried autoimmunity-associated terms in the enriched output and identified associated genes. After that, we retrieved interaction pairs using ToppCluster and CellChat database (Jin et al., 2020). In the end, we identified genes that are not only involved in autoimmunity, but have a mediator role in the immune signaling network. Using this approach, we observed several candidate pairs of genes, including TNFSF13B-TNSRSF13, IL10-IL10RA, IL21-IL21RA, IL6-IL6R, CXCL13-CXCR5, CXCL12-CXCR4, CCL21-CCR7, CCL19-CCR7 and CCL20-CCR6 in severe patients, which were enriched for autoimmune diseases, such as autoimmune thyroid diseases, lupus nephritis, autoimmune encephalomyelitis (Aust et al., 2004; Hirota et al., 2007; Kuwabara et al., 2009; Lee et al., 2010; Steinmetz et al., 2008). Candidate cytokine and chemokine ligand genes were expressed in various cell types in PBMC and BAL, including IL21 and CXCL13 from exhausted T cells of BAL, CXCL12 from mesenchymal cells, IL6 and CCL21 from endothelial cells, CCL19 from cDC and CCL20, TNFSF13B, and TNFSF13 from lung macrophages (**Figure 5E; Table S5**). These interaction pairs have been linked with auto-immunity (Klimatcheva et al., 2015; Wong et al., 2010). In addition, we analyzed single-cell studies (Arazi et al., 2019; Zhang et al., 2019) of rheumatoid arthritis and lupus nephritis patients and found that high expression levels of the candidate receptors in B cells and ligands in other cells were also observed, such as CXCL13 in helper T cells and CXCR5 in B cells in both studies (**Figure S15C, S15D**). However, more evidence is still required to infer the association between these interactions and autoimmunity in COVID-19 patients. Supported by the evidence above, we drew a network for potential mediator interactions of B cells and their associations with autoimmune disorders, where linkages with diseases, such as rheumatoid arthritis, systemic lupus erythematosus, were highlighted, as well as linkages with mouse phenotypes, such as abnormal immune tolerance and increased susceptibility to autoimmune disorder (**Figure 5F**). As a caveat, although using prior knowledge to prioritize gene and cell-associated functions and interactions may introduce biases, such approaches also have the potential to highlight key similarities and differences between different disease causes and clinical responses and improve our understanding of the molecular and cellular mechanisms at work.

### Functional Map and Immune Cell Interplay Landscape in COVID-19

As above, where highly significant enrichments of unique functions and pathways could be identified in the subtypes of multiple cell classes, such as neutrophils, platelets and B cells, we sought to get a more holistic understanding of COVID-19 specific cell class and subclass-level signatures, including T cell subtypes (**Figure S16 and S17**), we built an integrative functional map of all cell types in three compartments across multiple disease conditions using a highly integrated gene module set (**Figure 6A; Table S6**). All enriched functional associations in ToppCluster for gene modules of cell types and sub-clusters were depicted. They were grouped by disease conditions and compartments to show heterogeneity of cellular functions in different circumstances.

**Fig. 6.**
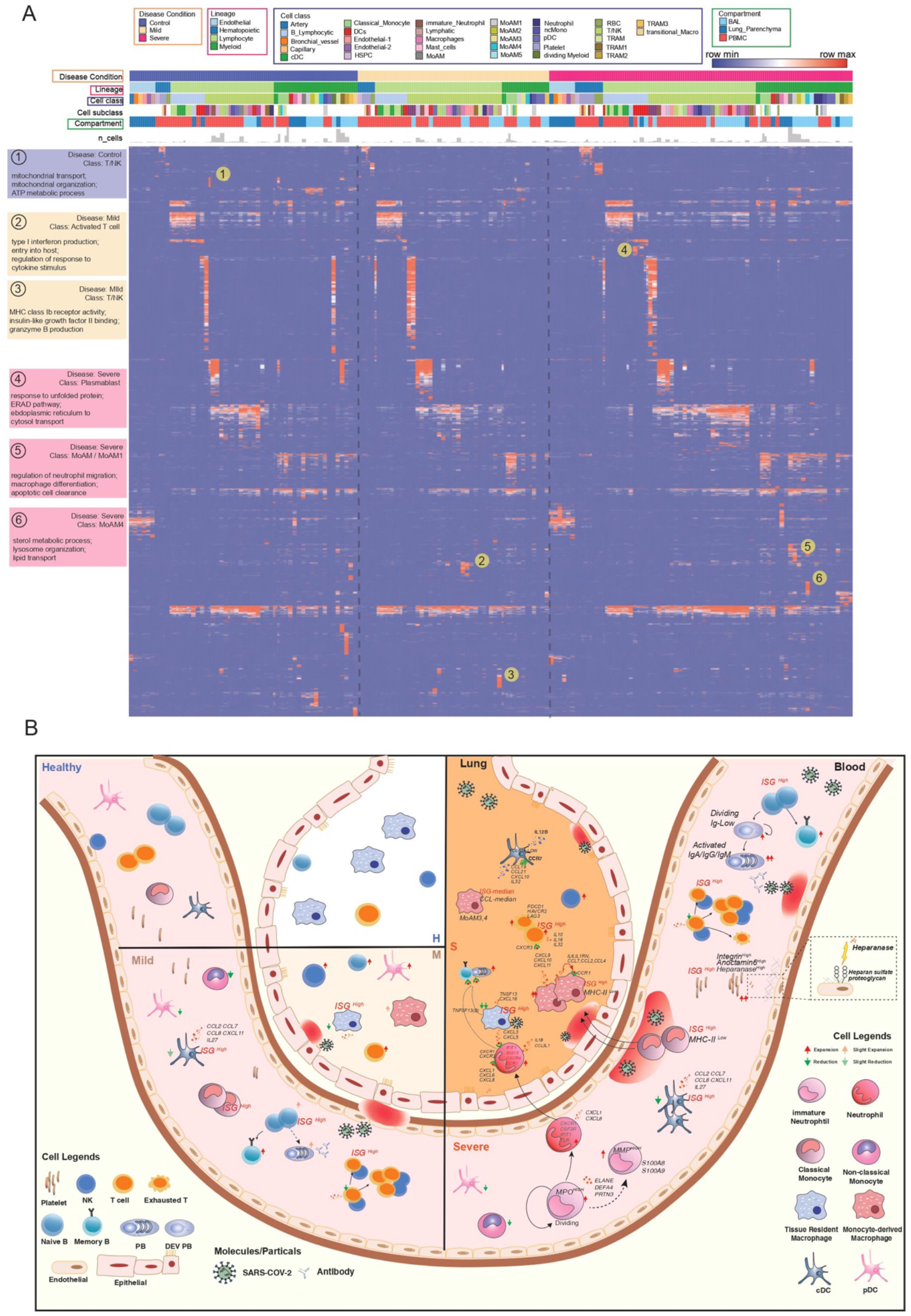
Comparative analysis of cell type specific gene signatures associated with lineage, class, subclass, compartment, and disease state in the COVID-19 atlas. (**A**) Enrichment scores of gene modules for all cell types across different compartments and COVID-19 conditions were generated by ToppCluster and shown on the heatmap. ToppCluster enriched functions from Gene Ontology, Human Phenotype, Mouse Phenotype, Pathway and Interaction databases were used to generate a feature matrix (cell types by features) and hierarchically clustered. Hot spots of the disease-specific enrichments were highlighted and details were shown on the left. More details can be found in Methods. (**B**) Summarizing predicted functions and interplay of immune cells in COVID-19 blood and lung. Aforementioned key observations in this study were shown in peripheral blood mononuclear cells (PBMC) and bronchoalveolar lavage (BAL) in healthy donors, mild and severe COVID-19 patients, including changes of cell abundance, specific marker genes, upregulated secretion, cell development and cell-cell interactions.

In the heatmap (**Figure 6A**), most enrichments were consistently observed across cells of healthy donors and COVID-19 patients. However, some unique patterns were also identified. For example, T cells and NK cells in healthy donors show enrichments of mitochondrial transport and ATP metabolic process, while activated T cells in mild patients show upregulation of type I interferon production and cytokine signaling. Enrichments of macrophage differentiation and neutrophil migration regulation were uniquely found in MoAM1 in severe patients (**Figure 6A**). The function map provides a high-level approach to investigate functional variations of cells across disease conditions and compartments. The predicted interplay of immune cells across multiple compartments and disease conditions is displayed in **Figure 6B**. Cell proportion changes, sub-cluster specific signatures and cell-cell interaction are also depicted.

### Similarity and Heterogeneity Between COVID-19 and Other Immune-mediated Diseases

To further analyze COVID-19 specific immune signatures, we compared immune cells from COVID-19 patients with cells in other immune-mediated diseases, including severe influenza (Lee et al., 2020), sepsis (Reyes et al., 2020) and multiple sclerosis (Schafflick et al., 2020). 404,125 cells were included after the integration of PBMC single-cell datasets (**Figure 7A, S18; Table S7**). Dynamic changes of cell abundance were compared in diseases versus healthy donors. Similar to COVID-19 patients, severe influenza patients also exhibited the reduction of non-classical monocytes, pDC, cDC and CD4+ TCM, but the effect of the former two types was smaller in magnitude (**Figure 7B**). However, the reduction of non-classical monocytes is more significant in severe COVID-19 patients than severe influenza or mild COVID-19 patients (**Figure 7B**). Notably, NK cell reduction is associated with COVID-19 severity, whereas T cell depletion is a more dramatic perturbation in severe influenza. Within these comparisons, the expansion of plasmablasts is consistently observed, whereas the accumulation of platelets is unique to SARS-CoV-2 and in particular, to severe COVID-19 clinical status (**Figure 7B**).

**Fig. 7.**
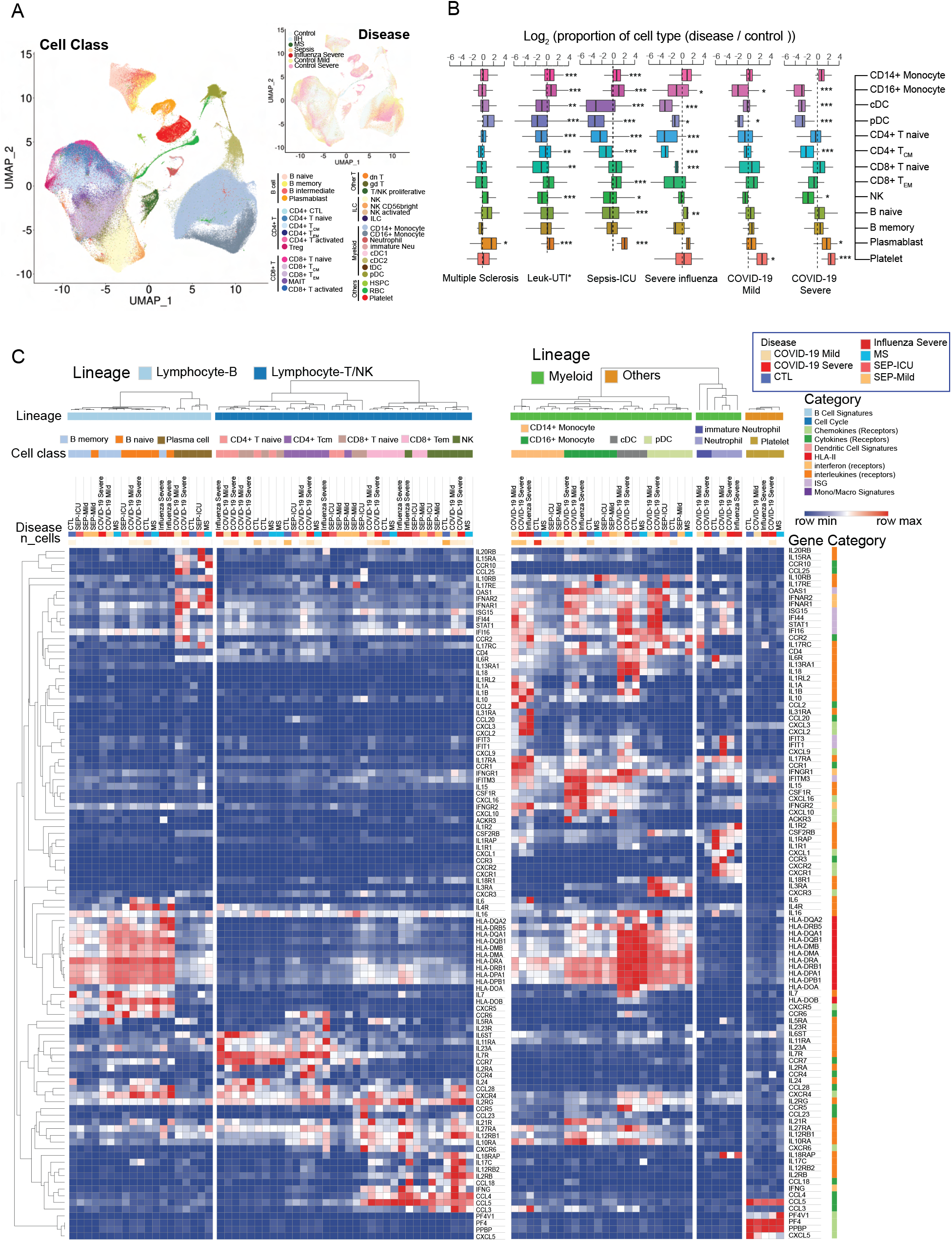
Comparative analysis of differentially-expressed immunoregulatory genes between COVID-19 and other immune-mediated diseases. (**A**) Uniform Manifold Approximation and Projection (UMAP) shows the distributions of cell types (Left) and diseases (Top right) after the integration of datasets in multiple studies. MS: multiple sclerosis; IIH: idiopathic intracranial hypertension. IIH patients were recruited as controls in the multiple sclerosis study. (**B**) Dynamic changes of immune cell types in different immune-mediated diseases compared to healthy controls. Log2(ratio) was calculated to show the levels of changes. *, p<0.05, **, p<0.01, ***, p<0.001. Statistical models can be found in the Methods. Leuk-UTI: sepsis patients that enrolled into UTI with leukocytosis (blood WBC ≥ 12,000 per mm3) but no organ dysfunction. (**C**) Normalized expression values of key genes involved in immune signaling and responses are shown for cell types across multiple diseases. Lowly expressed genes (maximal average expression level across all cell types in the heatmap is less than 0.5 after Log_2_CPM normalization) were removed.

In addition to dynamic changes of cell ratios, we also investigated the regulation of immune mediator genes across various diseases (**Figure 7C; Table S7**). IL-6 is an important factor of cytokine storms in COVID-19 (Zhao, 2020). As shown in the heatmap, naive B cells are the main sources of IL-6 in COVID-19 patients while CD14+ monocytes show the highest expression levels in severe influenza patients (**Figure 7C**). Specific ligands, including CXCL2, CXCL3, CCL20 were upregulated in both severe COVID-19 patients and severe influenza patients. CCR4 and IL2RA is uniquely high in CD4+ T cells of COVID-19 patients. Interestingly, most PBMC myeloid cell types displayed upregulated levels of interferon-stimulated genes in both COVID-19 and influenza, especially in COVID-19, where highest levels of ISGs in CD14+ Monocytes, cDC and pDC were observed.

## Discussion

In this work, we have constructed an innovative immune signature atlas of the blood and lung of COVID-19 patients using the integrated single cell RNA-sequencing data and Topp-toolkit. By virtue of systemic analysis of large sample size from multiple sampling sites, consistent immunopathology-associated changes of cell abundance and transcriptional profiles were observed in the circulating and lung immune repertoire of COVID-19 patients. The established single cell atlas and the provided public portal (https://toppcell.cchmc.org/) enables the query of candidate molecules and pathways in each of these processes.

Leveraging this approach, we identified three major candidate mechanisms capable of driving COVID-19 severity: (1) a cascade-like network of proinflammatory autocrine and paracrine ligand receptor interactions among subtypes of differentiating mononuclear, lymphoid, as well as other cell types; (2) the production of emergency platelets whose gene expression signatures implicate significantly elevated potential for adhesion, thrombosis, attenuated fibrinolysis, and potential to enhance the release of heparin-bound cytokines as well as further influence the activation of neutrophils causing further inflammatory cell recruitment and neutrophil netosis; and (3) the extrafollicular activation of naive and immature B cells via a multilineage network that includes monocytic subtypes and exhausted T cells of cytokines and interleukins with the potential to generate local antigen specific response to virus infected targets and collateral autoimmunity. More details will be discussed below.

We identified dramatically expanded macrophages which were marked by the loss of HLA class II genes and upregulation of interferon-stimulated genes. It implicates a key role for these activated macrophages involved in signaling network and less so in activation of adaptive T cell immunity. Among them, MoAM2 displayed hyperinflammatory responses and extraordinary high levels of signaling molecules, which are involved in both autocrine (e.g. IL-6, CCL2, CCL4 and CCL8) and paracrine (e.g. CXCL2, CXCL9, CXCL10 and CXCL11) signaling pathways. The former pathway contributed to the self-stimulation and development, which amplified the paracrine pathway for T cell and neutrophil chemoattraction. The latter two cell types in turn activated MoAMs with cytokines genes (CCL5, IL10 of T cells and IL1B of neutrophils, respectively). Based on the intercellular and multifactor complexity of the signaling cascade we have outlined, to effectively control a malignant inflammatory cascade, it may be essential to consider simultaneously targeting multiple nodes of this network of cytokines and interleukins. In addition, HLA-DR^low^ monocytes, likely reflecting dysfunctional cells, were observed in severe infection. This, along with evidence of emergency myelopoiesis with immature circulating neutrophils into the circulation was detected in severe COVID-19. These neutrophils had transcriptional programs suggestive of dysfunction and immunosuppression not seen in patients with mild COVID-19. As such, we have presented evidence for the contribution of defective monocyte activation and dysregulated myelopoiesis to severe COVID.

Platelet expansion is uniquely observed in COVID-19 versus other immune-mediated diseases.

Strikingly, these activated platelets were highlighted with abnormal thrombosis and upregulated heparanase, a procoagulant endoglycosidase that cleaves anti-coagulation heparan sulfate constituents on endothelial cells and potentially causes thrombotic vascular damages. Additionally, heparanase-cleaved heparan sulphate (HS) fragments were capable of stimulating the release of pro-inflammatory cytokines, such as IL1B, IL6, IL8, IL10 and TNF through the TLR-4 pathway in PBMC (Goodall et al., 2014), further contributing to the hyperinflammatory environment in COVID-19 patients. Since heparanase is recognized as a hallmark in tumor progression and metastasis (Jayatilleke and Hulett, 2020), we hypothesize COVID-19 infection could be associated with higher occurrence of lung tumor metastasis. However, more data is required to support it. Pro-neutrophil secreted proteins (e.g. ELANE, DEF4) of neutrophil extracellular trap (NET), which have been reported to be associated with higher risk of morbid thrombotic events (Zuo et al., 2021). Approaches to combatting NETs could a potential anticoagulation treatment (Thålin et al., 2019).

We propose a signaling network which potentially shapes the differentiation of B cells towards the formation of autoantibodies. Proliferation and activation of inflammatory myeloid cells and the formation of exhausted CD4+ T helper around an area of direct or indirect viral tissue injury leads to the production of a set of interleukins and cytokines known to have both direct cell activating and maturing effects on naïve and immature B cells. Previous report had revealed the exaggerated extrafollicular B cell response, which is part of a mechanism that stimulates somatic mutation and maturation of B cells to produce plasma cells with specificity for antigens present in the vicinity of tissue damage sites (Farris and Guthridge, 2020). In the absence of macrophages or dendritic cells to restrict self vs non-self, the presence of IL-10, IL-21, CXCL13 CXC10, IL-6 and others acting on receptors present in naïve and immature B cells leads to the selection and maturation of self-reactive maturation of B cells clones with formation of autoantibodies. Many of these COVID-19-activated genes (e.g. CXCL13, CCL19, CCL20, TNFRSF13) are known to be genetically associated with rheumatoid arthritis, lupus, and risk of developing autoimmune disease in humans and mouse models. The development of different patterns of autoimmunity may be the main hallmark of “Long Haul” Covid disease and could explain why some individuals develop different autoantibodies and suffer different forms of clinical consequences depending on which antigens drive the B-cell maturation. Thus, an additional prediction that could be made based on these findings and our network model is that among individuals treated with corticosteroids at the time these auto-immunogenic processes are activated, there should be a protective effect and lower likelihood of developing post acute sequela of Covid.

Consistent and varied compositional changes and gene patterns of immune cells were identified in COVID-19, influenza and sepsis. Expansion of plasmablasts, as well as the reduction of non-classical monocytes, are more significant changes in severe COVID-19 patients, while the depletion of T cells is more dramatic in severe influenza patients. The accumulation is a unique immune hallmark of COVID-19 within the selected diseases, which contributes to the coagulation abnormalities and thrombosis, a key cause of fatality in COVID-19 patients. Different signaling gene patterns were identified across immune-mediated diseases, with CCR4 only highly expressed in CD4+ T cells of COVID-19 patients, which might be related with extravasation of these cells (Spoerl et al., 2021). Upregulated interferon-stimulated genes of myeloid cells in PBMC revealed the inflammatory environment of COVID-19.

Collectively, using the COVID-19 single cell atlas data exploration environment, we have illustrated is that researchers are now enabled to systematically explore, learn, and formulate new hypotheses within and between compartments, cell types, and biological processes, and provided access to these reprocessed datasets through a suite of explorative and evaluative tools. Moreover, we have shown different hypotheses can be developed and explored using the approaches that we have outlined and the database that we have provided. Certainly additional critical information will also be obtained using approaches that include in situ spatial, temporal data as well as those of viral products and viral and inflammatory-process affected complexes. Next steps for improving its ability to be mined more deeply will be based on additional statistical methods that extend the current ToppCell / ToppGene Suite based on fuzzy measure similarity, Page-Rank, and cell-cell signaling approaches.

There are several limitations in our study. Different studies used various standards of COVID-19 severity definition. To generalize conclusions, we simplified disease conditions into several universal groups. Prospectively, a standardized definition of disease stages will assist to the accuracy of future studies. Additionally, the timing of sample collection was not considered as a variable in this study, rather disease stages were used to consolidate data across samples. W lack follow-up data of patients with sequela, which will be helpful for understanding the long-haul effects of the disease.

## Acknowledgements

We thank Pablo Garcia-Nieto, Ambrose Carr and Jonah Cool and the Chan Zuckerberg Initiative for hosting the data on cellxgene. We acknowledge suggestions and help from Greta Beekhuis. Some figures were created using https://biorender.com. Funding for this study was provided by LungMap (U24 and HL148865), Digestive Health Center (P30, DK078392) and Harold C. Schott Foundation funding of the Harold C. Schott Endowed Chair, UC College of Medicine. We thank the support from Pediatric Cell Atlas and high performance computational cluster of CCHMC.

## Author Contributions

Conceptualization, K.J. and B.A.; Methodology, K.J., B.A., E.B., A.M. and J.Y.W; Investigation, K.J. and B.A.; Writing – Original Draft, K.J.; Writing – Review & Editing, K.J., B.A., M.E.R., A.M., D.A.P.K, S.S.G. and S.B.; Funding Acquisition, B.A.; Resources, K.J.; Data Curation, K.J. and E.B.; Visualization, E.B.; Supervision, B.A..

## Declaration of Interests

The authors declare no competing interests.

## Resource availability

### Lead contact

Further information and requests for resources should be directed to and will be fulfilled by the Lead Contact, Bruce Aronow.

### Materials Availability

This study did not generate new unique reagents.

### Data and Code Availability

Public single-cell RNA-seq datasets of PBMC in COVID-19 patients are available on NCBI Gene Expression Omnibus and European Genome-phenome Archive, including <u>GSE150728</u>, <u>GSE155673</u>, <u>GSE150861</u>, <u>GSE149689</u> and EGAS00001004571 (or <u>FastGenomics</u>). BAL single-cell RNA-seq datasets of COVID-19 patients are available on <u>GSE145926</u> and <u>GSE155249</u>. Lung Parenchyma single-cell RNA-seq data are available on <u>GSE158127</u>.

Single-cell RNA-seq data of sepsis patients are available on the Single Cell Portal <u>SCP548</u> and <u>SCP550</u>. Data of multiple sclerosis patients are available on <u>GSE128266</u>. Data of severe influenza patients are available on <u>GSE149689</u>.

Gene modules of all datasets analyzed using ToppCell web portal are available on COVID-19 Atlas in <u>ToppCell</u>, including gene modules from either a single dataset or an integrated dataset. Gene modules from the integration of specific cell types, such as B cells and neutrophils are also listed in ToppCell. More details are listed in Figure1A and Table S1. An interactive interface of integrated PBMC data and subclusters of immune cells will be public on <u>cellxgene</u>.

Codes of preprocessing, normalization, clustering and plotting of single-cell datasets will be available on <u>github</u>.

## Methods

### Single-cell RNA-seq data source

To have a comprehensive understanding of immune cells in different repertoires, we collected 8 public COVID-19 single-cell RNA-seq datasets of multiple compartments, including peripheral blood mononuclear cells, bronchoalveolar lavage and lung biopsy, which in total covered over 43 healthy donors, 22 mild/moderate, 42 severe and 2 convalescent COVID-19 patients. More details can be found in Figure 1A and Table S1. Lung biopsy samples were taken from the explanted lung or post-mortem lungs of COVID-19 patients(Bharat et al., 2020). Various criteria were used in these publications to describe COVID-19 severity. For example, we found asymptomatic, mild, moderate and floor COVID-19 patients under the definition of non-severe COVID-19 patients in our data sources. A recent paper used the WHO score of COVID-19 severity to categorize disease conditions of patients(Wilk et al., 2020b), which is a more standardized and robust approach for the description of disease stages. However, in order to address the issue of missing information for disease stratification and to simplify the comparison, we grouped disease conditions into three groups, including healthy donors, mild COVID-19 patients and severe COVID-19 patients. Convalescent patients were excluded in some of our analysis for simplification. Sequencing data of healthy donors in Guo et al. was excluded since it was not from the same institute(Guo et al., 2020).

We also collected PBMC single-cell RNA-seq data from 29 sepsis patients(Reyes et al., 2020) and 4 multiple sclerosis(Schafflick et al., 2020) patients for comparative analysis of immune-mediated diseases (**Figure 1A; Table S1**). Data sources can be found in Data Availability.

### Data preprocessing and normalization

For datasets with raw UMI counts, we first removed cells with less than 300 detected genes or less than 600 UMI counts. Then cells with more than 15% counts of mitochondrial genes were filtered out. Genes expressed in less than 5 cells were removed. After quality control, we finally harvested 483,765 high-quality cells from 8 studies (**Table S1**). We normalized the total UMI counts per gene to 1 million (CPM) and applied log_2_(CPM+1) transformation for heatmap visualization and downstream differential gene expression analysis. Steps above were done in Scanpy(Wolf et al., 2018).

For some datasets that only provide processed and normalized *h5ad* or *rds* files, we checked their preprocessing procedures in the original publications and confirmed that stringent quality control procedures were used. Most of them used the default normalization approach in the Seurat or Scanpy pipeline. We transferred them to log_2_(CPM+1) to make data consistently normalized. We also prepared corresponding raw count files for data integration.

### Integration of PBMC datasets and BAL datasets using Reciprocal PCA in Seurat

We input raw count files of 5 preprocessed PBMC datasets into Seurat and created a list of Seurat objects. Reciprocal PCA procedure (https://satijalab.org/seurat/v3.2/integration.html~reciprocal-pca) was used for data integration. First, normalization and variable feature detection were applied for each dataset in the list. Then we used *SelectIntegrationFeatures* to select features for downstream integration. Next, we scaled data and ran the principal component analysis with selected features using *ScaleData* and *RunPCA*. Then we found integration anchors and integrated data using *FindIntegrationAchnors* and *IntegrateData*. RPCA was used as the reduction method. After integration, we scaled data and ran PCA on integrated expression values. UMAP was generated using the top 30 reduced dimensions with *RunUMAP*. The same approach was also used in BAL data integration and multi-disease integration. We also used it for the integration of specific cell types across multiple datasets, for example, the integration of neutrophils from PBMC and BAL datasets. Compared with standard workflow and SCTransform (https://satijalab.org/seurat/v3.2/integration.html) in Seurat, we found Reciprocal PCA is much less computation-intensive and time-consuming, making the integration of multiple large single-cell datasets feasible.

### Cell Annotations using canonical markers after unsupervised clustering

Cell annotations were assigned in each dataset and then mapped to the integrated data. For some datasets without available cell annotations, we first used unsupervised clustering in Scanpy. Detailed steps include (1) detecting top 3,000 highly variable genes using *pp.highly_variable_genes*; (2) scaling each gene to unit variance on highly variable genes using *pp.scale*; (3) running PCA using *arpack* approach in *tl.pca*; (4) finding neighbors using *pp.neighbors*; (5) running leiden clustering with resolution of 1 using *tl.leiden* (resolutions were determined swiftly based on the size and complexity of data). More details can be found in the code (point to it). For datasets with available annotations, we checked their validity and corrected wrong annotations. For example, hematopoietic stem and progenitor cells (HSPC) were mistakenly annotated as “SC&Eosinophil” in the original paper(Wilk et al., 2020a) and were corrected in our annotation.

After unsupervised clustering, well recognized immune cell markers were used to annotate clusters, including CD4+ T cell markers such as TRAC, CD3D, CD3E, CD3G, CD4; CD8+ T cell markers such as CD8A, CD8B, NKG7; NK cell markers such as NKG7, GNLY, KLRD1; B cell markers such as CD19, MS4A1, CD79A; plasmablast markers such as MZB1, XBP1; monocyte markers such as S100A8, S100A9, CST3, CD14; conventional dendritic cell markers such as XCR1, plasmacytoid dendritic cell markers such as TCF4; megakaryocyte/platelet marker PPBP; red blood cell markers HBA1, HBA2; HSPC marker CD34. Exhaustion-associated markers, including PDCD1, HAVCR2, CTLA4 and LAG3 were used to identify exhausted T cells.

Additionally, other markers were used for annotations of lung-specific cells, including AGER, MSLN for AT1 cells; SFTPC, SFTPB for AT2 cells; SCGB3A2, SCGB1A1 for Club cells; TPPP3, FOXJ1 for Ciliated cells; KRT5 for Basal cells; CFTR for Ionocytes; FABP4, CD68 for tissue-resident macrophages; FCN1 for monocyte-derived macrophages, TPSB2 for Mast cells. More details can be found in Table S2.

### Cell Annotations using Azimuth

To better annotate T cells in our study, we applied Azimuth (https://satijalab.org/azimuth/), a tool for reference-based single-cell analysis developed in Seurat version 4.0(Hao et al., 2020). High-quality PBMC single-cell data in Azimuth was used as the reference for label projection. After removing annotations with low prediction scores or low mapping scores, we got a collection of well-annotated T cell subtypes, including CD4+ Cytotoxic T cell, CD4+ Naive T cell, CD4+ Central Memory T cell, CD8+ Naive T cell, CD8+ Effector Memory cell, gamma-delta T cell, double-negative T cell. CD4+ Effector Memory T cell and CD8+ Central Memory T cell were found by Azimuth but removed later because of low scores. Apart from annotations of T cell subtypes, we also found CD56-bright NK cell, intermediate B cell and Memory B cell using Azimuth.

### Sub-clustering for specific cell types

Sub-clustering was used for the discovery of subtypes or distinct stages of a specific cell type. In our work, we applied sub-cluster for various immune cell types, including classical monocytes, neutrophils, conventional dendritic, B cells and platelets. First, all cells in the specific cell type were integrated using the same procedure as PBMC data integration. Then Louvain clustering (resolution = 0.5, except for sub-clustering of classical monocytes where resolution = 0.3) was applied to detect sub-clusters of those cells. Importantly, neutrophils, cDCs and B cells were retrieved from both PBMC and BAL, whereas classical monocytes and platelets were only retrieved from PBMC.

### Generation of ToppCell gene modules

ToppCell (https://toppcell.cchmc.org/) was designed to parallelly analyze transcriptional profiles of single-cell datasets by organizing differential expressed gene modules in a customized hierarchical order. In our study, we hierarchically annotated cells with multiple layers, including compartments, disease conditions, lineages, cell classes and sub-clusters. All the cells were grouped into specific hierarchical categories. For example, “PBMC_severe COVID-19_myeloid cells_classical-monocytes_cMono1” represents cells belonging to cMono1 (a sub-cluster of classical monocytes) in PBMC of severe COVID-19 patients. With hierarchically ordered cell annotations, we calculated their DEGs in a hierarchical way as well. We defined customized ranges for comparisons and applied t-test based on normalized expression values. More details can be seen on ToppCell website. Usually, the top 200 most differentially genes in each comparison were picked up as the gene modules for the selected cell group, which are the starting point of downstream analysis, including gene enrichment in ToppGene and interaction inference in ToppCluster. All gene modules in our study were curated in COVID-19 Atlas (https://toppcell.cchmc.org/biosystems/go/index3/COVID-19Atlas) and ImmuneMap (https://toppcell.cchmc.org/biosystems/go/index3/ImmuneMap) on the ToppCell website.

### Gene Enrichment Analysis using ToppGene

Abundant gene modules were generated with ToppCell. After that, we used ToppGene (https://toppgene.cchmc.org/) for gene enrichment analysis. Genes in each gene module were sent to ToppGene platform as input for enrichment in different domains. GO-Molecular Function, GO-Biological Process and GO-Cellular Component and Mouse Phenotype were usually used for enrichment. P values of enrichment results were adjusted using the Benjamini-Hochberg procedure.

### Generation of Functional Association Heatmap using ToppCluster

Genes in gene modules of selected cell types or sub-clusters were sent to ToppCluster (https://toppcluster.cchmc.org/). Then multi-group functional enrichment was drawn for input gene modules and -log_10_(adjusted p-value) was used as the gene enrichment score to represent the strength of association between gene modules and pathways. Scores greater than 10 were trimmed to 10. Pathways from Gene Ontologies, including Molecular Functions, Biological Process and Cellular Component in the option list were used for the enrichment of gene modules in myeloid cells, B cells and platelets. In order to gain a broader knowledge of immunothrombosis-related pathways, “Pathway” and “Mouse Phenotype’’ in the option list were also selected for enrichment. Morpheus was used for visualization of the heatmap (https://software.broadinstitute.org/morpheus/).

### Cell Interaction Inference in immunothrombosis activities and cytokine signaling pathways

CellChat was used to infer the signaling network in the BAL of severe patients (**Figure S8B**). All 3 categories of interactions were used in the database *CellChatDB.human*. Over-expressed ligands or receptors in each cell type were first identified for further identification of over-expressed interaction pairs. Then cytokine, chemokine and IL signaling probability between multiple cell types was inferred using *computeCommunProb* and *computeCommunProbPathway*.

ToppCell was used to infer interactions in immunothrombosis. We first selected genes related to coagulation or immunothrombosis pathways from subtypes of endothelial cells, platelets, neutrophils, classical monocytes and monocyte-derived macrophages by filtering the output of ToppCluster (**Figure S12A**). Then we used CellChatDB as the knowledge base to find the subset of genes participating in cell-cell interaction, including genes involved in signaling via secretion, cell-cell contact and extracellular matrix interaction. These genes in each cluster were sent to ToppCluster to infer the interaction network using protein-protein interactions (PPI) between those genes.

### Statistics of Cell Proportion Changes in Different Disease Stages

Cell proportion differences between disease groups for specific types and subtypes (**Figure 2, Figure S2-S4**) shown on box plots were measured by Mann-Whitney test (Wilcoxon, paired=False). Significance between two disease conditions were shown on the top.

To investigate the dynamic changes of cell proportions across various immune-mediated diseases, we followed the approach in recent literature (Lee et al., 2020) (**Figure 7B**). For each disease condition, we computed the relative ratio of each cell type in individual disease samples divided by individual healthy samples. Log_2_ transformed values were shown in the box plot. Then we calculated relative ratios of each cell type between all sample pairs of healthy donors as a control. To compute the significance, we used a two-sided Kolmogorov-Smirnov (KS) test using relative ratios in diseases and those values in healthy donors.

### Generation of Volcano Plots

We first calculated differential expressed genes using *tl.rank_genes_groups* in Scanpy. Adjusted p values and log fold changes in the output were used as the input of volcano plots. R package *EnhancedVolcano (Blighe et al., 2018)* was used to draw figures.

### Construction of COVID-19 Functional Enrichment Map

In order to characterize functional properties of cell types and subtypes observed in BAL, PBMC, and lung parenchymal samples from control, mild, and severe COVID-19 patient samples, we used the library of gene expression signatures (“Gene Module Report” from <u>ToppCell</u>) as an input to the ToppCluster enrichment analyzer web server (Kaimal et al 2010). Using categories of *Gene Ontology*, *Human Phenotype*, *Mouse Phenotype*, *Pathway* and *Protein Interaction,* a matrix was constructed using minus log P enrichment values for each celltype gene list and then all cells and enriched features could be clustered and ordered based on their shared or distinct properties that could then be associated with lineage, cell subclass, tissue compartment, and disease state.

## Supplementary Information

Table S1. Metadata of data sources and generated resources in this study, relative to Figure 1.

Table S2. Severity-associated cell distribution in multiple compartments of COVID-19 patients, relative to Figure 2.

Table S3. Gene modules and DEGs of neutrophil and macrophage sub-clusters, relative to Figure 3.

Table S4. Gene modules and pro-thrombosis genes in platelets of COVID-19 patients, relative to Figure 4.

Table S5. Developing plasmablast signatures and autoimmunity-associated signatures of COVID-19 B cells, relative to Figure 5.

Table S6. Enrichment scores of COVID-19 Functional Map, relative to Figure 6.

Table S7. Dynamic changes of abundances and signatures across immune-mediated diseases, relative to Figure 7.

---

**Figure S1.**
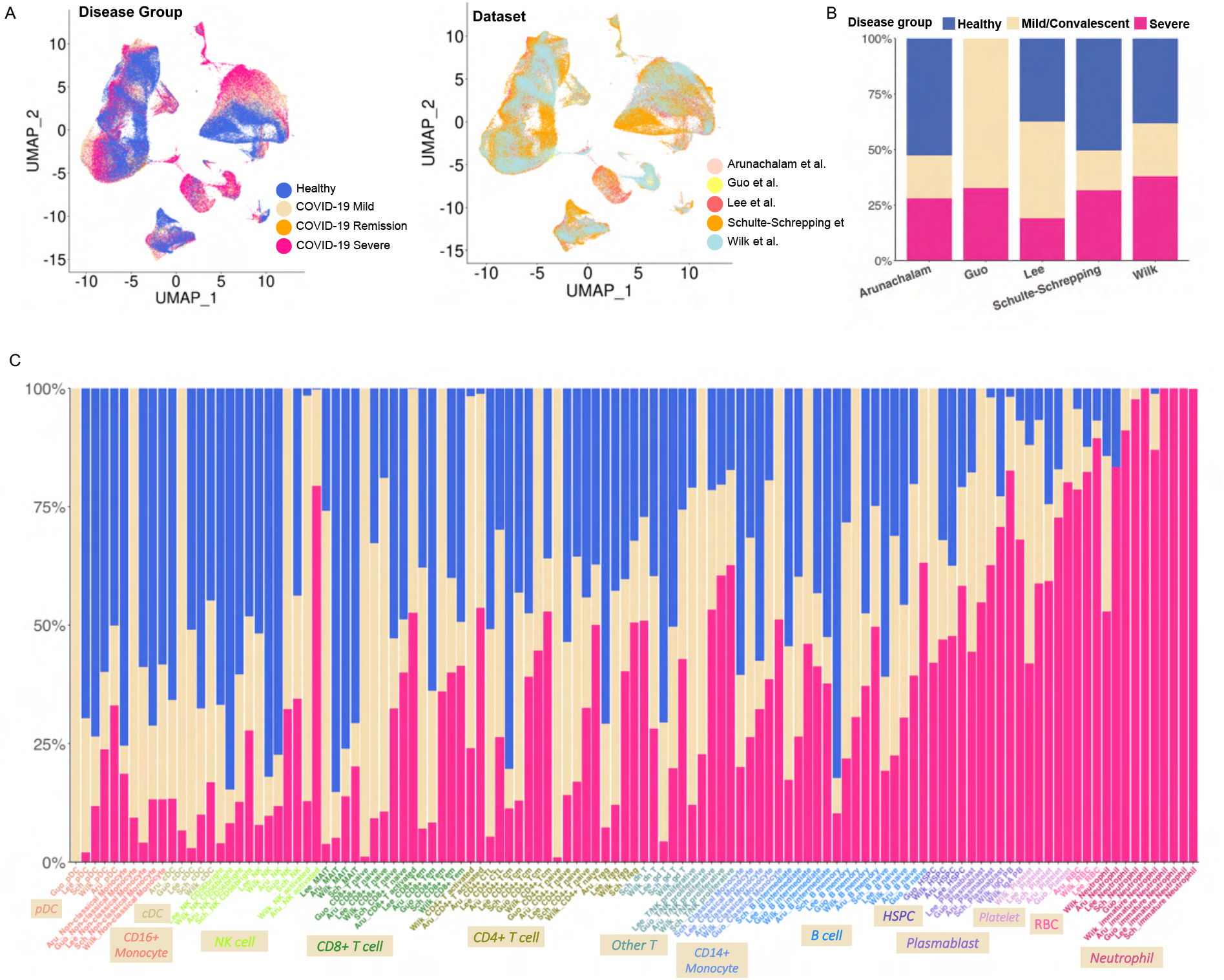
Cell distribution and abundance in the integrated COVID-19 PBMC data, relative to Figure 2. (**A**) Distributions of COVID-19 conditions (Left) and data sources (Right) for the integrated PBMC data are shown on the same UMAP of Figure 2A. (**B**) Bar plot depicts distributions of disease conditions in 5 individual PBMC single-cell datasets. Percentages of 3 disease conditions in each dataset is shown on y axis. (**C**) The integrated bar plot shows percentages of 3 disease conditions in each cell type per dataset. Dataset abbreviations and cell types were concatenated to show disease distributions of specific cell types in the selected datasets. These labels are colored by their cell type designations and ordered by the ascending percentages of COVID-19 conditions.

**Figure S2.**
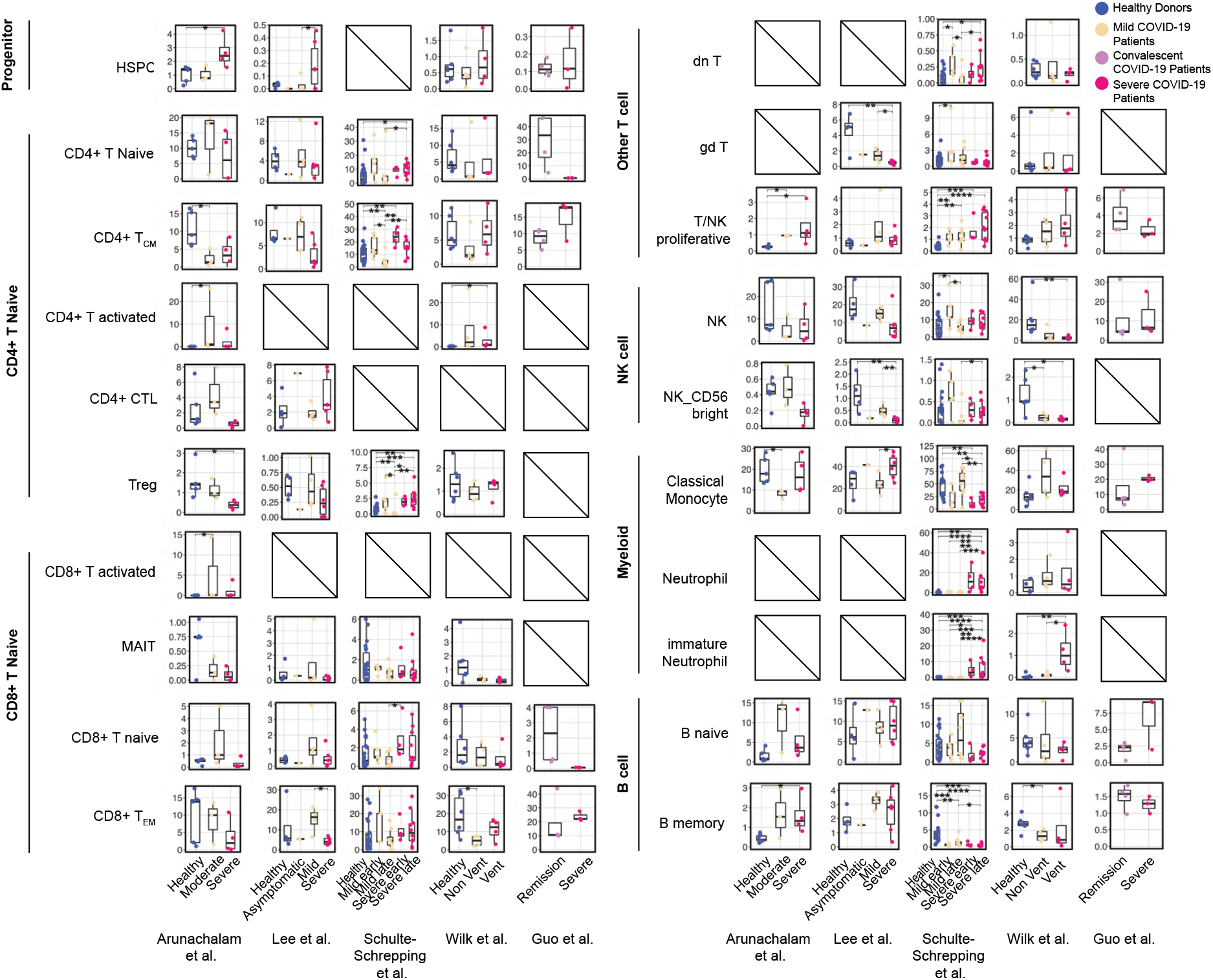
Dynamic changes of cell type abundances in five COVID-19 PBMC datasets, relative to Figure 2. Relative abundances and differences of major cell types in each single cell dataset are shown and compared to controls per each disease condition, per each single-cell dataset. Box plots of all cell types in PBMC are shown except for the 5 highlighted cell types shown in Figure 2B. Statistical methods are the same with Figure 2B.

**Figure S3.**
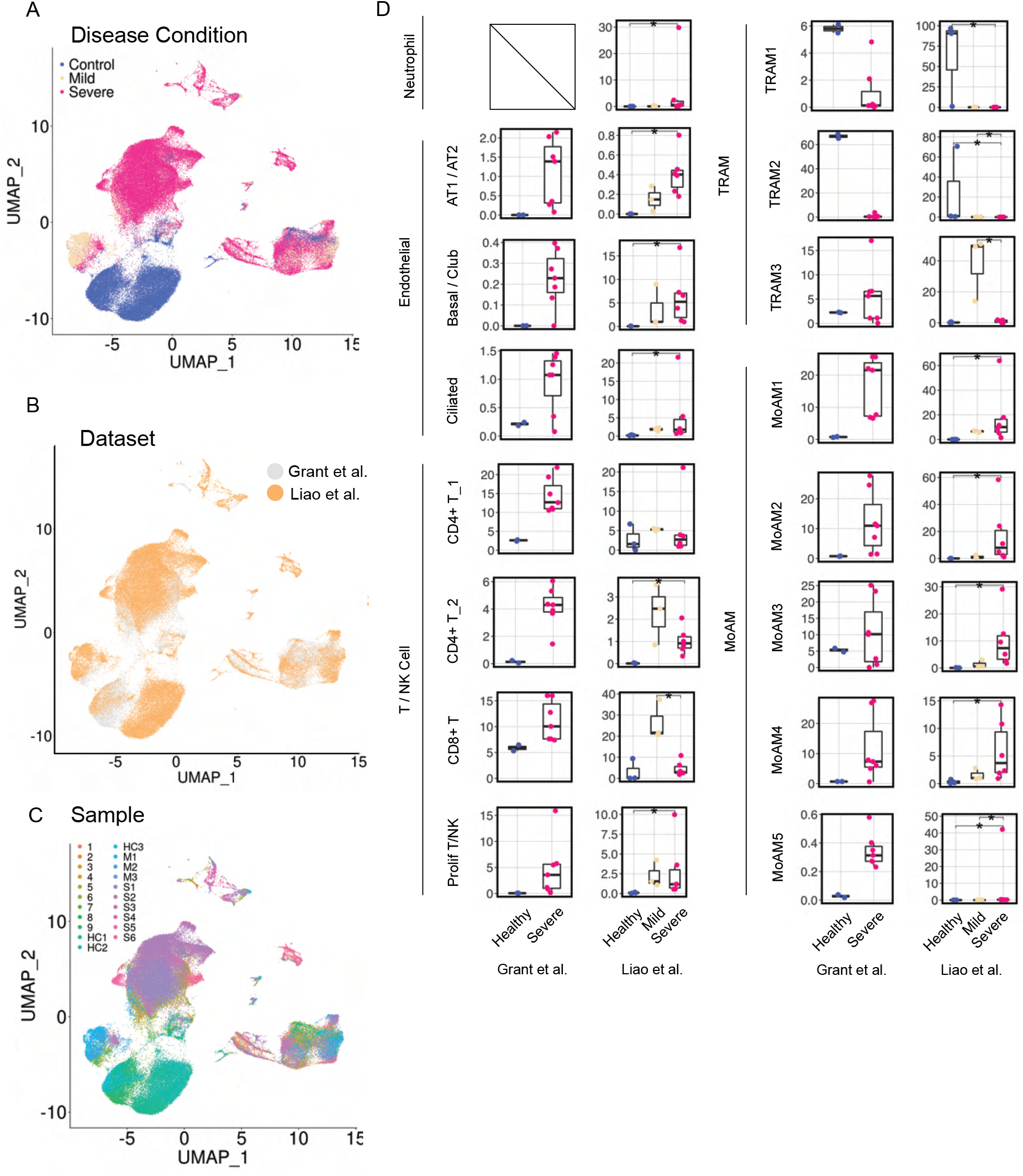
Cell distributions and dynamic changes in the integrated COVID-19 BAL data, relative to Figure 2. (**A-C**) Distributions of disease conditions (A), data sources (B) and samples (C) are shown on the same UMAP of Figure 2C. (**D**) Box plots depict dynamic changes of cell types across COVID-19 conditions in BAL that are not covered in Figure 2D. Statistical methods are the same with Figure 2B.

**Figure S4.**
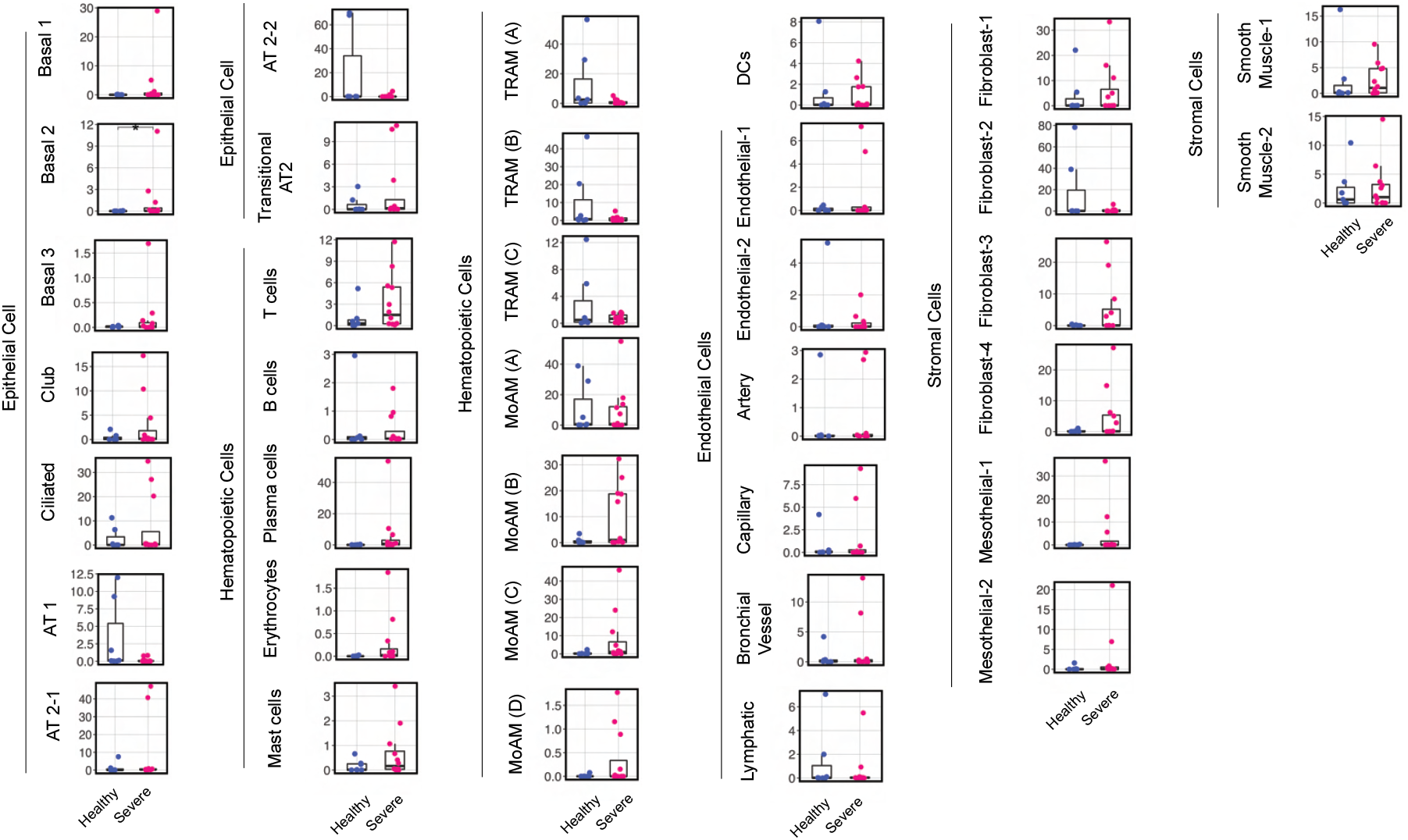
Cell type abundance changes in COVID-19 lung parenchyma dataset, relative to Figure 2. Box plots depict percentages of cell types in control samples and severe COVID-19 samples. We used cell type clusters identified in the original publication but modified cell naming of macrophage subtypes to distinguish monocyte derived macrophage subtypes present in BAL fluid samples. Statistical methods are the same with Figure 2B.

**Figure S5.**
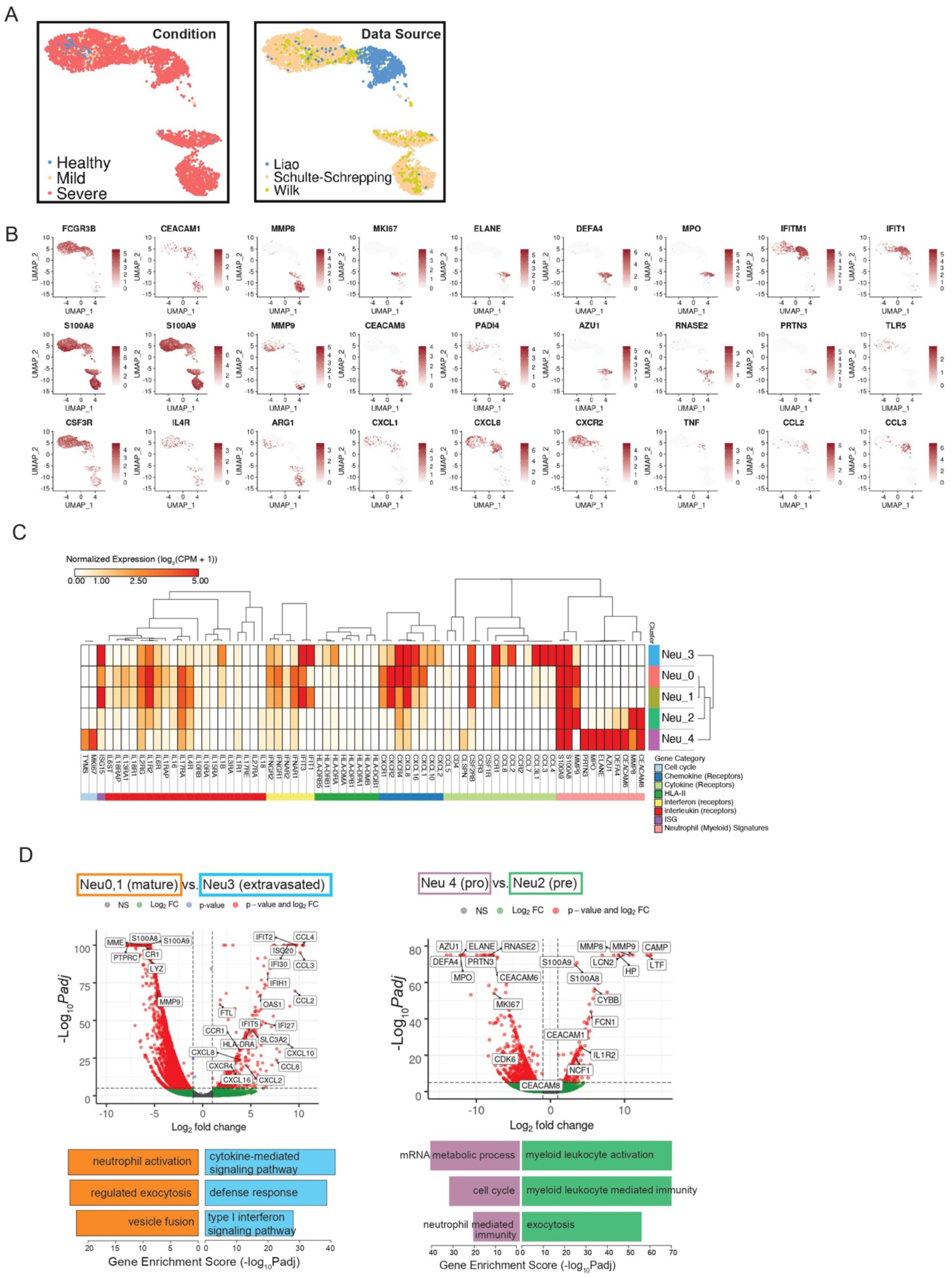
Sub-cluster-specific genes of neutrophils of COVID-19 patients, relative to Figure 3. (**A**) Distribution of disease conditions (Left) and data sources (Right) for the integrated neutrophil data on the same UMAP of Figure 3A. (**B**) UMAPs of neutrophil sub-cluster-associated genes from Figure 3C. Normalized expression values for each gene were used. (**C**) Normalized expression values of neutrophil-associated genes and other important immune signatures are shown for 5 neutrophil sub-clusters. Lowly expressed genes (genes with maximal average expression level across all neutrophil sub-clusters less than 0.5 after Log_2_CPM normalization) were removed from the gene pool of cytokines, chemokines, ISGs, interleukins, interferons, corresponding receptors and MHC-II. (**D**) The volcano plot depicts differentially expressed genes between circulating mature neutrophils (Neu0,1) and extravasated neutrophils (Neu3) (Left); as well as DEGs between pro-neutrophils (Neu4) and pre-neutrophils (Neu2) (Right). Statistical methods are the same with Figure 5C. Representative enriched biological processes (Gene Ontology) are shown in the bottom.

**Figure S6.**
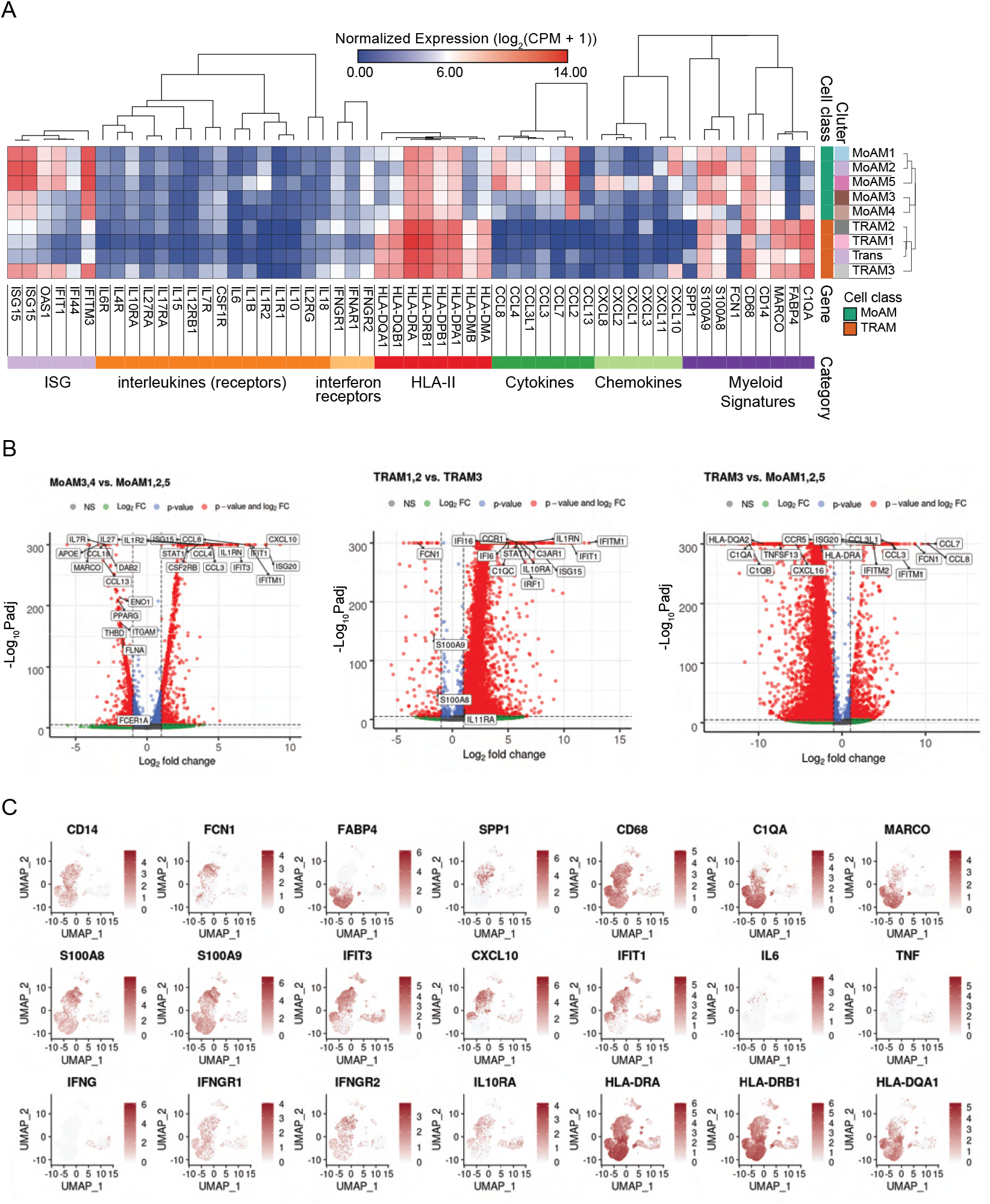
Macrophage-related signatures in the integrated BAL data, relative to Figure 3. (**A**) Normalized expression values of myeloid-cell-associated genes and other important immune signatures are shown for 9 macrophage sub-clusters. Lowly expressed genes (genes with maximal average expression level across all macrophage sub-clusters less than 0.5 after Log_2_CPM normalization) were removed from the gene pool of MHC-II, cytokines, chemokines, ISGs, interleukins, interferons and their receptors. (**B**) Volcano plots were drawn for DEGs of MoAM3,4 versus MoAM1,2,5 (Left) and TRAM 1,2 versus TRAM3 (Middle) and TRAM3 versus MoAM1,2,5 (Right). Statistical methods are the same with Figure 5C. (**C**) Normalized expression values were shown on the same UMAP of Figure 3B for important genes including macrophage signatures, ISGs, interferons, receptors and MHC-II.

**Figure S7.**
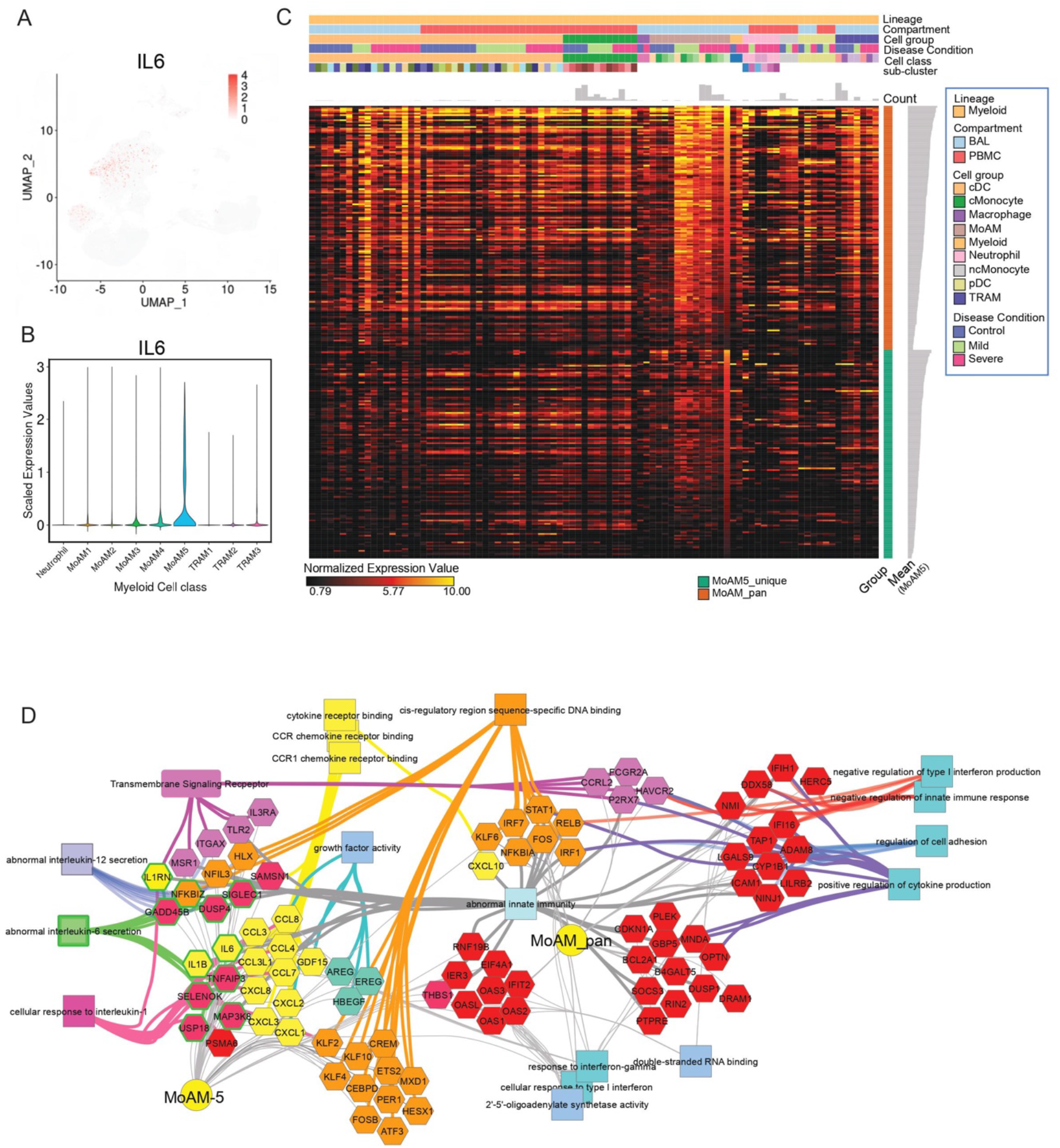
A uniquely-activated monocyte-derived cell type (MoAM5) exhibits a broad signature of cytokines, chemokines, and interleukins including IL6, relative to Figure 3. (**A**) Normalized expression values of IL6 on the same reference UMAP of integrated BAL data as Figure 3B. (**B**) Scale expression levels of IL6 for each macrophage sub-cluster on the violin plot. (**C**) Heatmap of expression levels of pan-MoAM signatures and MoAM5-specifc signatures in all myeloid cells in both PBMC and BAL. (**D**) Network of functional and phenotypic associated pan-MoAM signatures and MoAM5-specific signatures from (C). Associations were retrieved from ToppGene enrichment results. IL6 is highlighted in the network. As a caveat, the MoAM5 subtype represented a small fraction among the BAL MoAM subtypes and the majority of these cells were observed in a single severely-affected individual.

**Figure S8.**
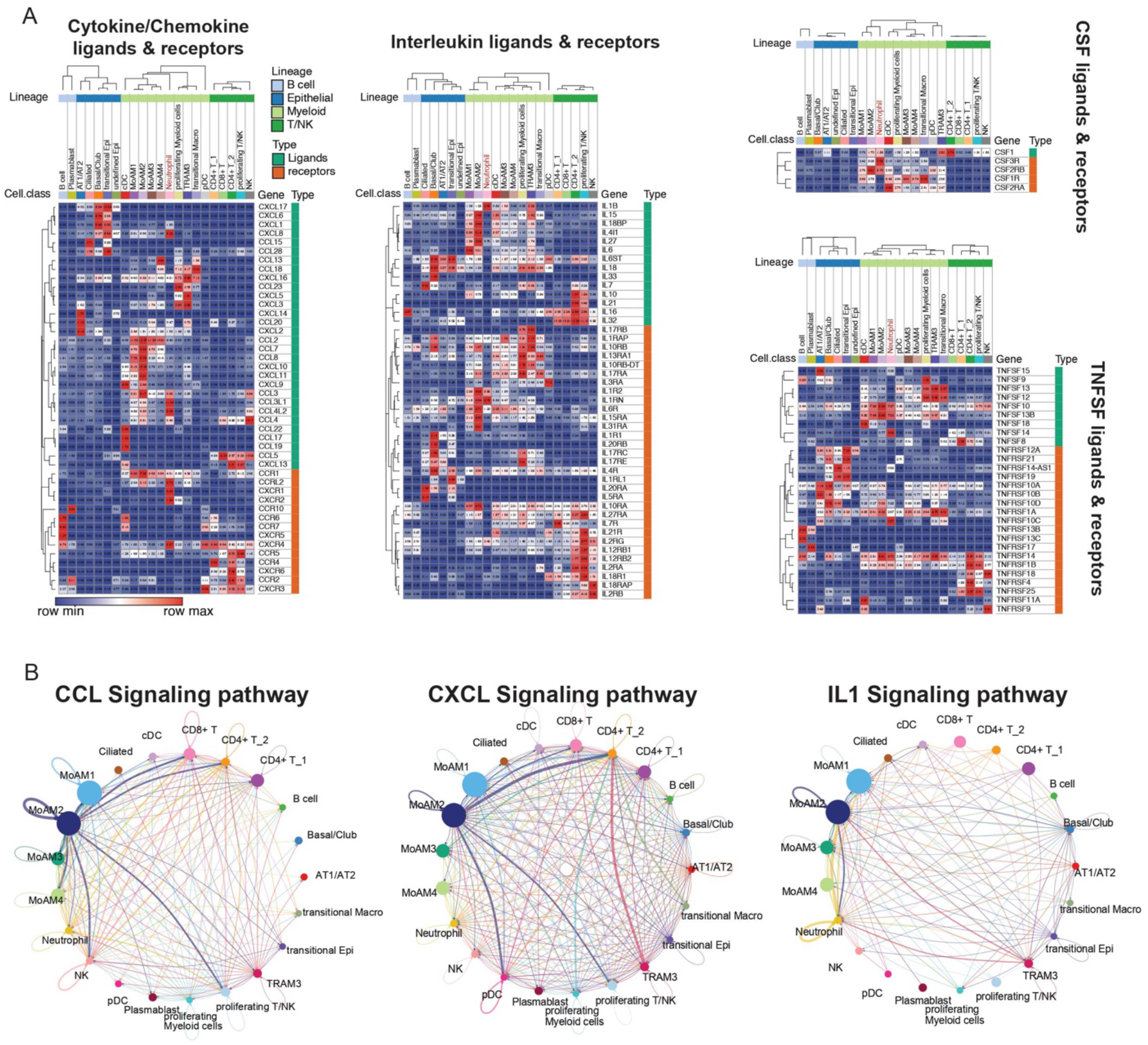
Cell type and cell subtype-specific divisions of cytokine, chemokine, and interleukin signaling pathways in BAL of severe COVID-19 patients, relative to Figure 3. (**A**) Heatmap of expression patterns of ligands and receptors in cytokine, chemokine, interleukin, CSF and TNFSF signaling pathways across cell types of BAL in severe patients. Average normalized expression values were shown and lowly expressed ligands or receptors (maximal normalized expression value for a row in the heatmap < 0.5) were removed. To reduce bias, MoAM5 was removed because cells in the cluster were mainly from one patient. Cell types that have less than 5% cells from severe patients were removed, including TRAM1 and TRAM2. Neutrophils are highlighted in the heatmap. (**B**) Interaction network of BAL cells in severe patients using CellChat. CCL, CXCL and IL1 signaling pathways were shown. The width of edges represents the strength of interactions and the size of nodes represents the abundance of cell types.

**Figure S9.**
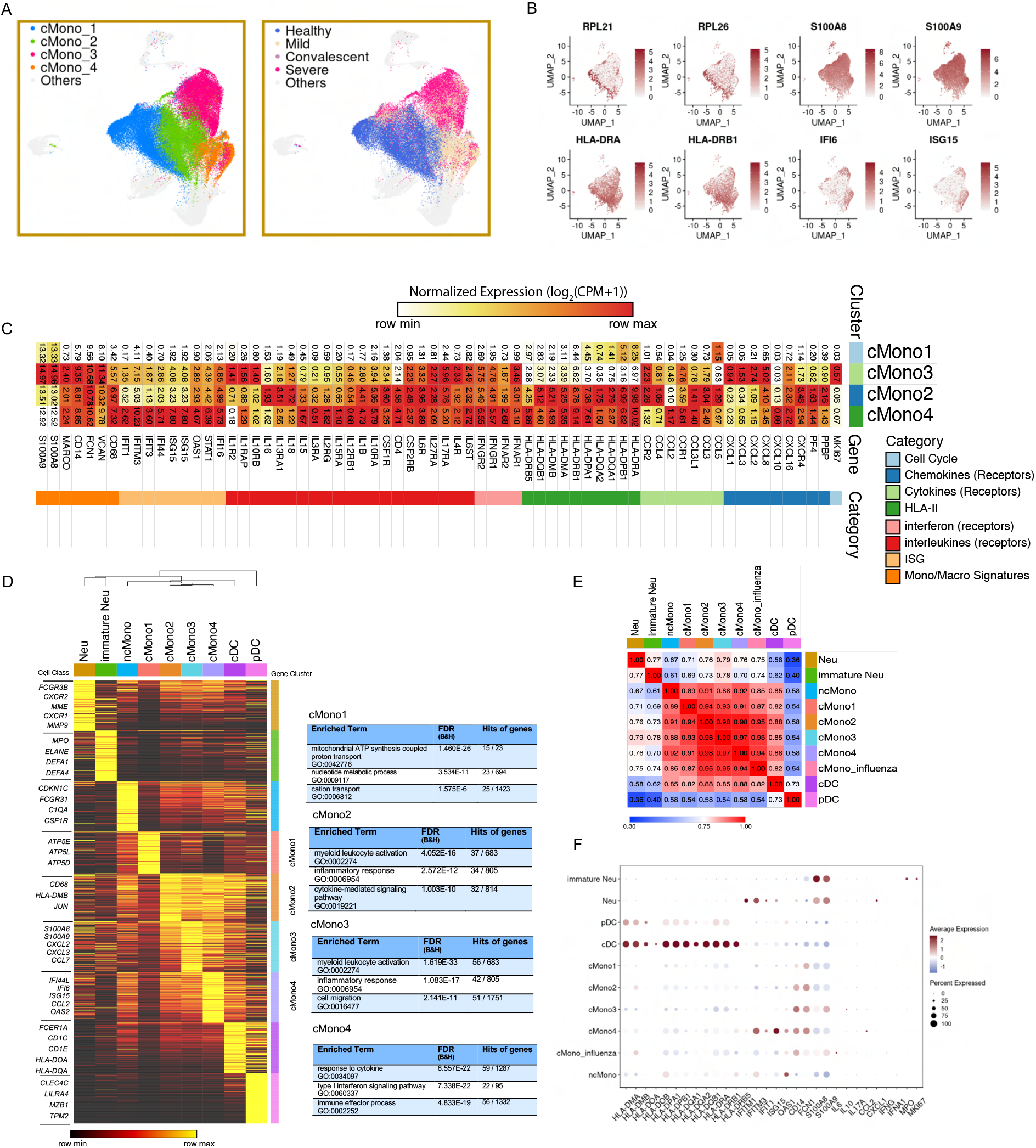
Characteristics of sub-clusters of classical monocytes in the integrated COVID-19 PBMC data, relative to Figure 3. (**A**) UMAPs of 4 sub-clusters (Left) and COVID-19 conditions (Right) of classical monocytes are shown. Grey dots are other myeloid cells in the UMAP of integrated PBMC myeloid data. (**B**) UMAPs of normalized expression values of specific signatures for classical monocyte sub-clusters. (**C**) Normalized expression values of monocyte-associated genes and other important immune signatures are shown for 4 classical monocyte sub-clusters. (**D**) Gene modules of classical monocyte sub-clusters, as well as other myeloid cell types in the integrated PBMC myeloid data. Representative genes in each module are shown on the left. ToppGene enrichment results for classical monocyte sub-clusters are shown on the right. Columns are clustered using hierarchical clustering. (**E**) Similarity matrix of myeloid cell types using genes in (D). Pearson correlation was used to evaluate similarity. (**F**) Dot plot of MHC-II, ISGs, interleukin genes and cell cycle genes for each myeloid cell type. Scale values were used.

**Figure S10.**
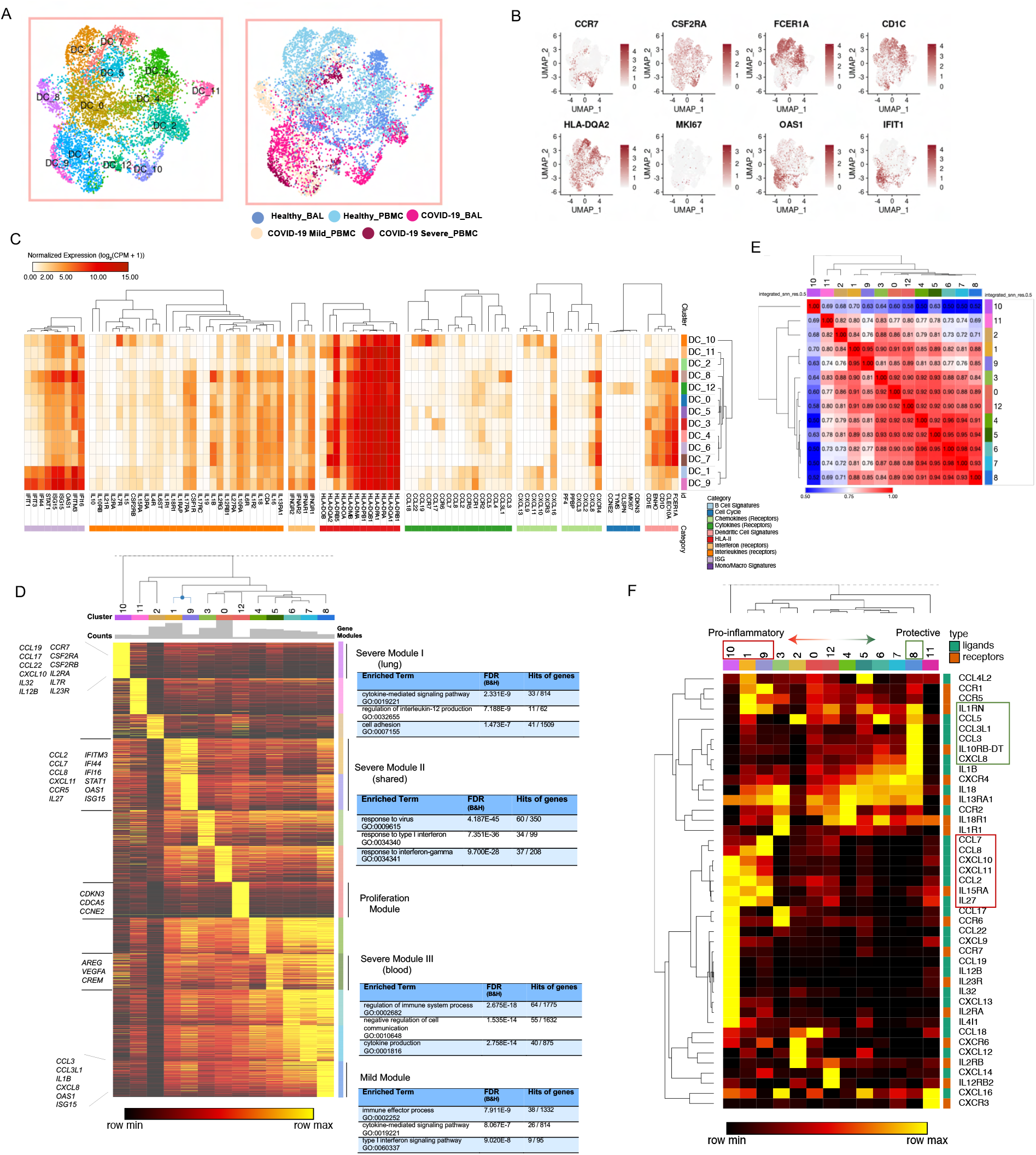
Features of conventional dendritic cell sub-clusters and polarized signaling genes, relative to Figure 3. (**A**) UMAPs of 13 sub-clusters (Left) and sources (Right) of conventional dendritic cells after data integration. (**B**) Normalized expression values of sub-cluster-specific genes on the UMAP. (**C**) Normalized expression values of cDC-associated genes and other important immune signatures are shown for 13 cDC sub-clusters. (**D**) Gene modules of cDC sub-clusters with 200 most significantly upregulated genes in each module. Representative genes are shown on the left. Gene enrichment results of some modules from ToppGene are shown on the right. (**E**) Similarity matrix of sub-clusters using genes in (D). Pearson correlation was used for similarity scores and hierarchical clustering was applied for rows and columns. (**F**) The heatmap shows the clustering of signaling genes, including cytokines, chemokines, interleukins and their receptors. Red boxes highlight severe patients associated sub-clusters and their upregulated genes. Green boxes highlight mild patients-associated sub-clusters and their upregulated genes.

**Figure S11.**
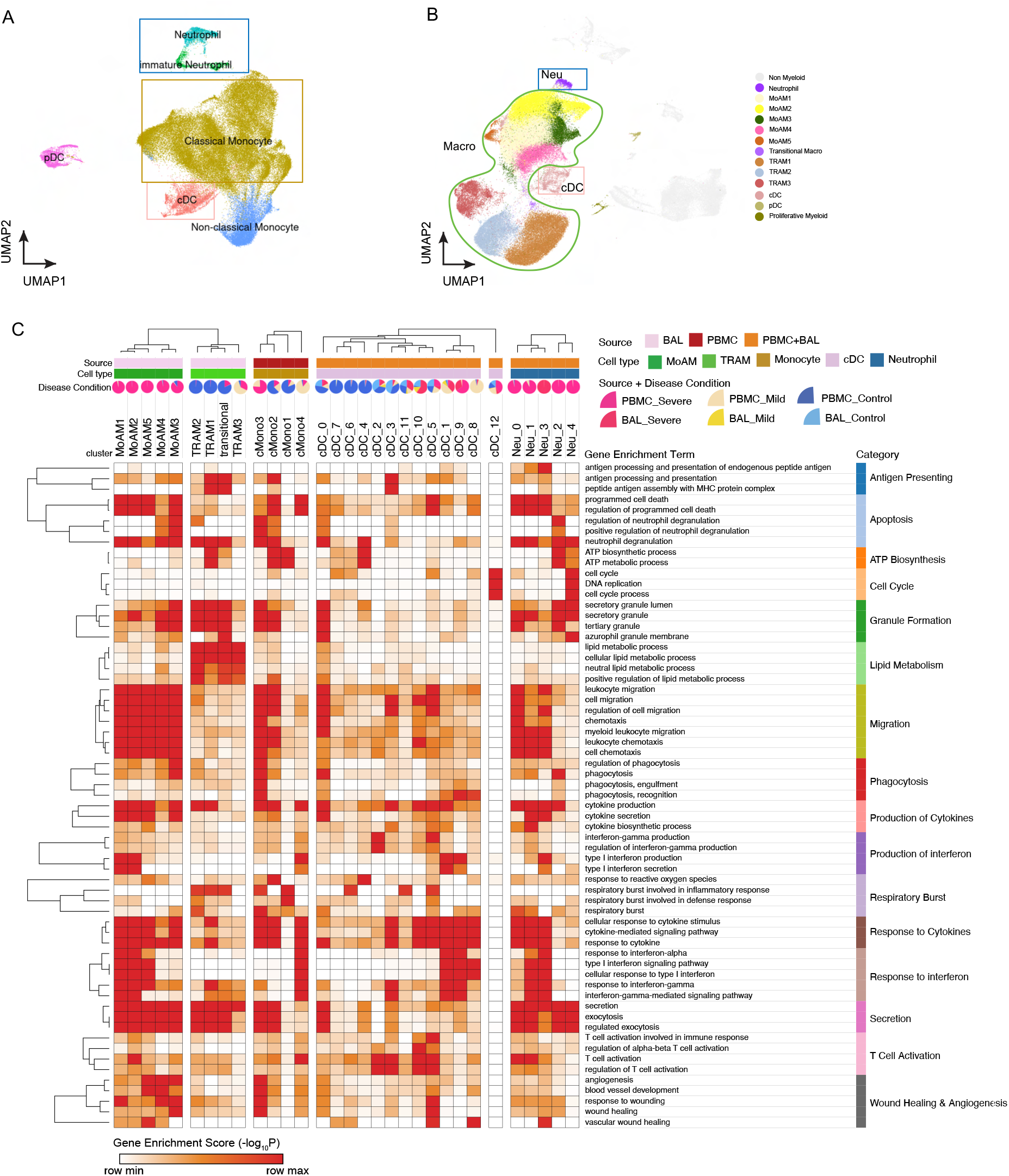
Landscape of myeloid cells in the integrated PBMC and BAL data, relative to Figure 3. (**A-B**) UMAPs of myeloid cells in integrated PBMC (A) and BAL (B) data. Cell types which were further clustered are highlighted in different colors. (**C**) The heatmap shows associations between subclusters of myeloid cells and myeloid-cell-associated pathways, such as antigen presenting, T cell activation, phagocytosis etc. Gene enrichment scores, defined as -log_10_(adjusted p value), were calculated as the strength of associations. Pie charts showed the proportions of COVID-19 conditions in each sub-cluster.

**Figure S12.**
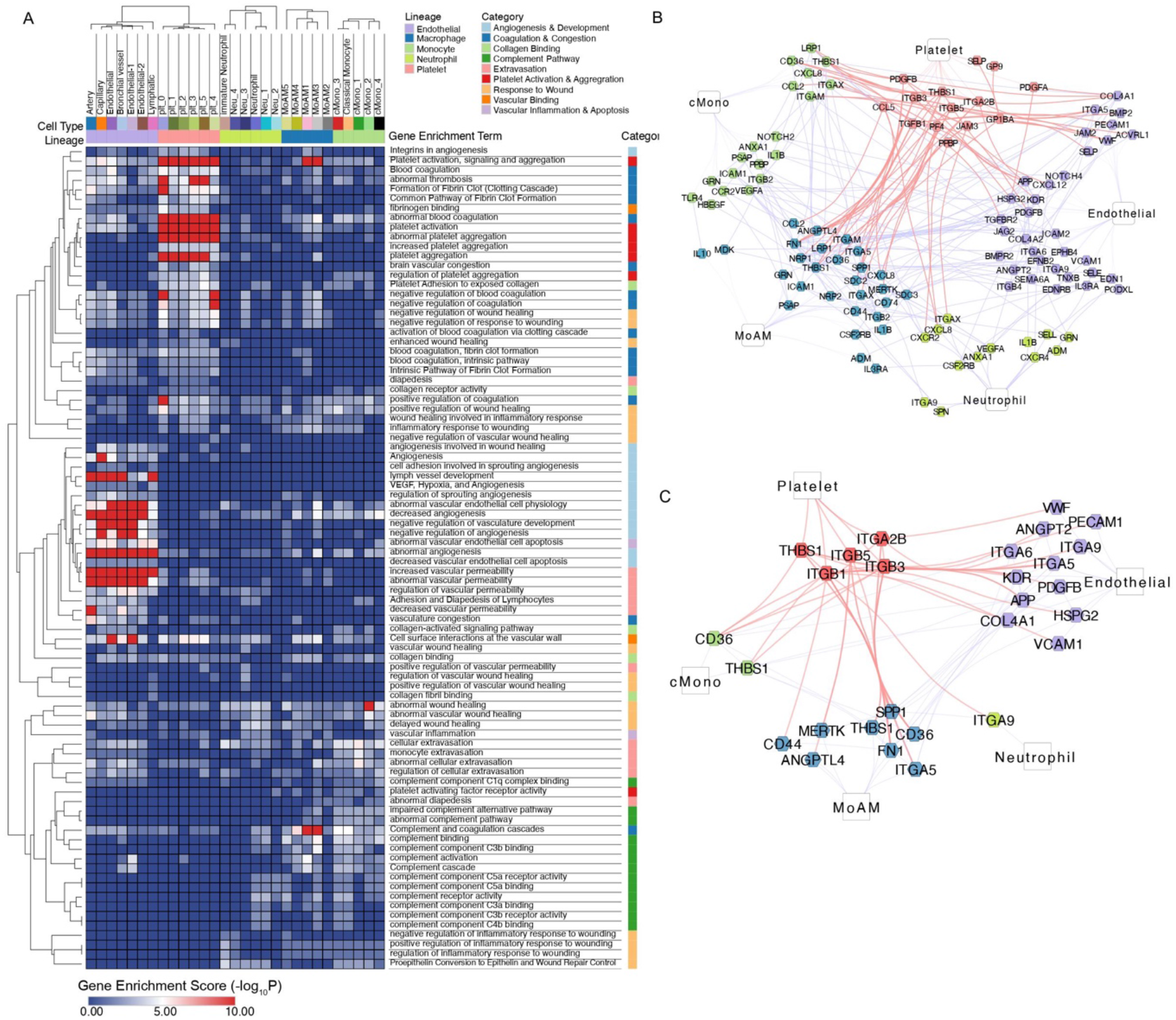
Gene expression signatures of cell types and subtypes activated by COVID-19 are extensively associated with coagulation, hemostasis, and thrombosis-associated pathways, functions, and knockout phenotypes, relative to Figure 4. (**A**) Functional association heatmap of gene signatures from COVID-19 cell types demonstrates differential enrichment for pathways associated with coagulation, vascular permeability, complement, extravasation, platelet activation and aggregation, response to wounding, as shown. Gene modules of cell types and sub-clusters that participate in these pathways were used to calculate enrichment scores. (**B**) Network of upregulated genes in coagulation/thrombosis-associated pathways (A) shows the potential gene-gene interactions in immunothrombosis of COVID-19 patients. CellChat and ToppCell/ToppGene protein-protein ligand receptor and cell adhesion interaction databases were used to find interaction pairs among upregulated genes. (**C**) A new network derived from (B) shows integrin-associated interactions between platelets and other cells.

**Figure S13.**
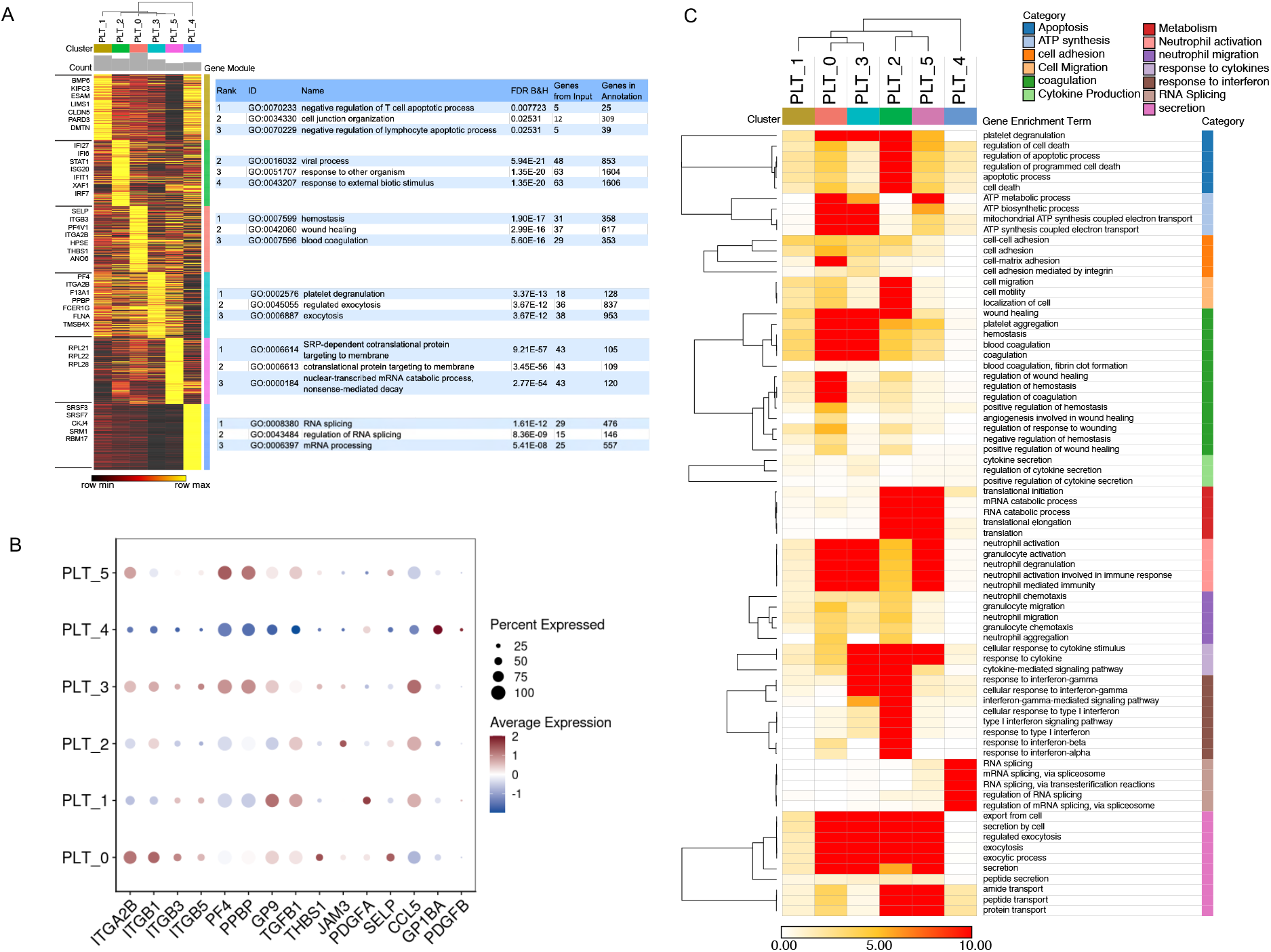
Emergence of platelet subtypes implicating functionally significant alternative roles in hemostasis, coagulation, wound response, and neutrophil recruitment and activation, relative to Figure 4. (**A**) The heatmap shows ToppCell gene modules of 6 platelet sub-clusters in COVID-19 PBMC. Each gene module contains 200 most significant genes for each sub-cluster and important genes are shown on the left. Gene enrichment analysis was conducted using ToppGene and top enrichment results from biological processes (Gene Ontology) are shown on the right. (**B**) Dot plot of integrin and other platelet-associated genes. Scale values are shown on the figure. (**C**) Heatmap of associations between subclusters of platelets and platelet-associated pathways (Gene Ontology). Gene enrichment scores, defined as - log_10_(adjusted p value), were calculated and shown.

**Figure S14.**
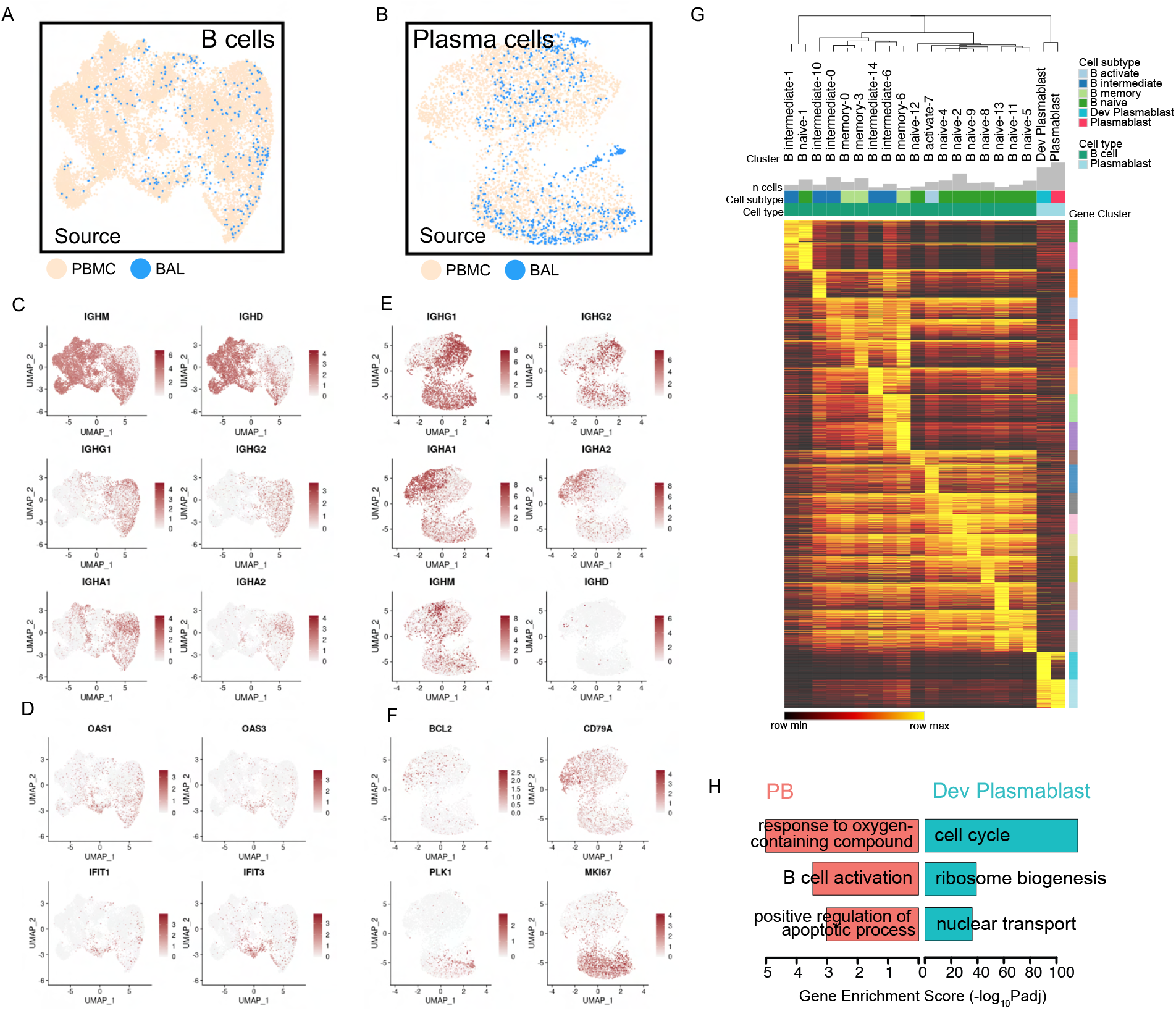
Consistent emergence of a series of early and maturing B cells and plasmablasts in BAL fluid and PBMC across multiple datasets, relative to Figure 5. (**A-B**) UMAPs of B cells (A) and plasmablasts (B) from multiple datasets. (**C-D**) UMAP of normalized expression values of immunoglobulin genes (C) and ISGs (D) for B cells. (**E-F**) UMAP of normalized expression values of immunoglobulin genes (E) and sub-cluster associated genes, such as cell cycle genes and B cell markers (F) for plasmablasts. (**G**) Gene modules of B cell sub-clusters and plasmablast subtypes with 200 most significant genes in each module. Hierarchical clustering was applied for columns. (**H**) Three representative enriched biological processes (Gene Ontology) are shown for these two subtypes using DEGs of plasmablasts in Figure 5C.

**Figure S15.**
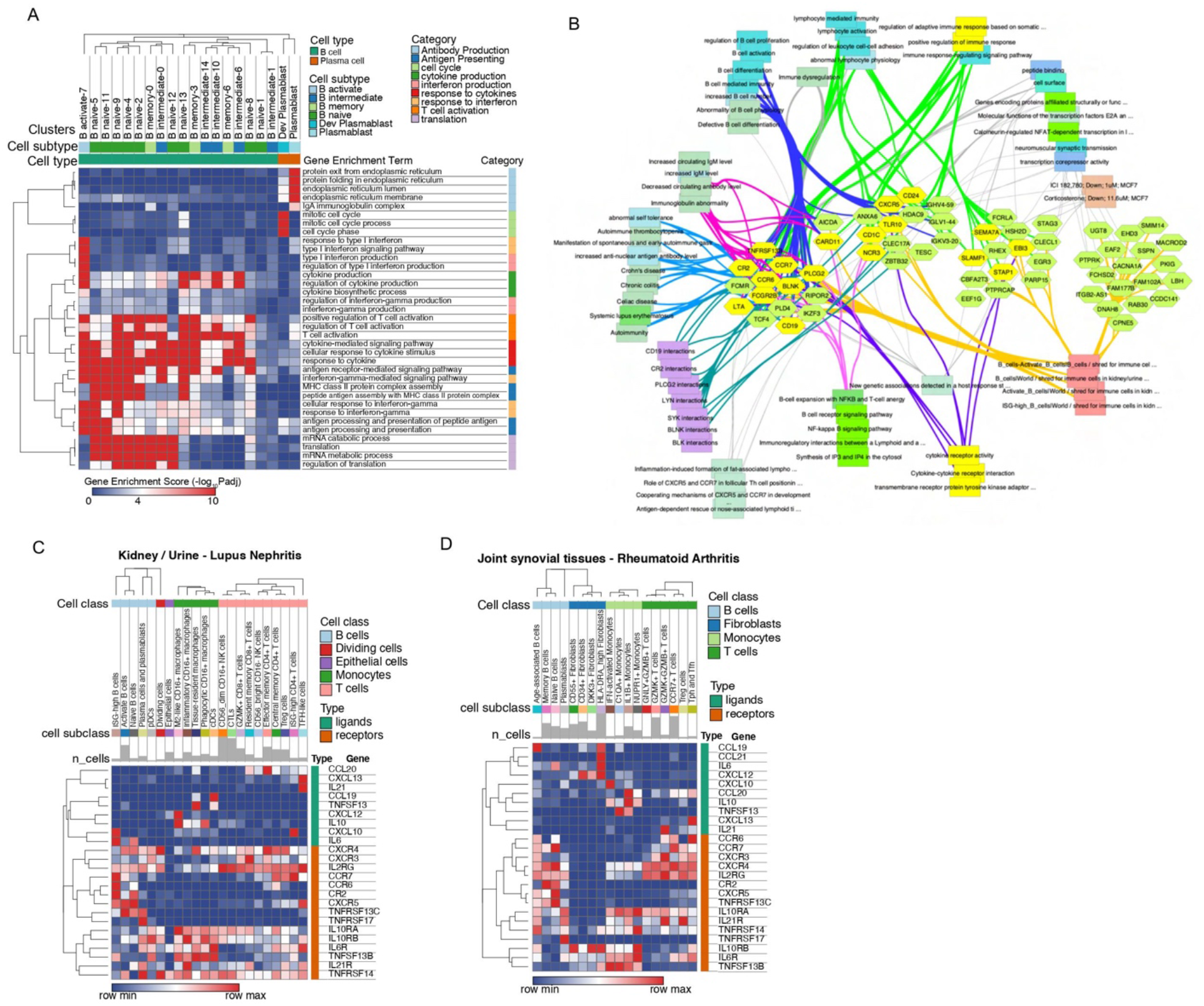
Gene Enrichment analysis of B cell subtypes and autoimmune-associated signatures, relative to Figure 5. (**A**) Heatmap shows gene enrichment scores of B-cell-associated pathways for each B cell sub-cluster and plasmablast subtype. (**B**) Pathway and function association network of upregulated genes in B cells of BAL in mild COVID-19 patients. (**C-D**) Heatmaps show normalized expression levels of autoimmune-associated ligands and receptors (Figure 5E) in lupus nephritis (C) and rheumatoid arthritis (D).

**Figure S16.**
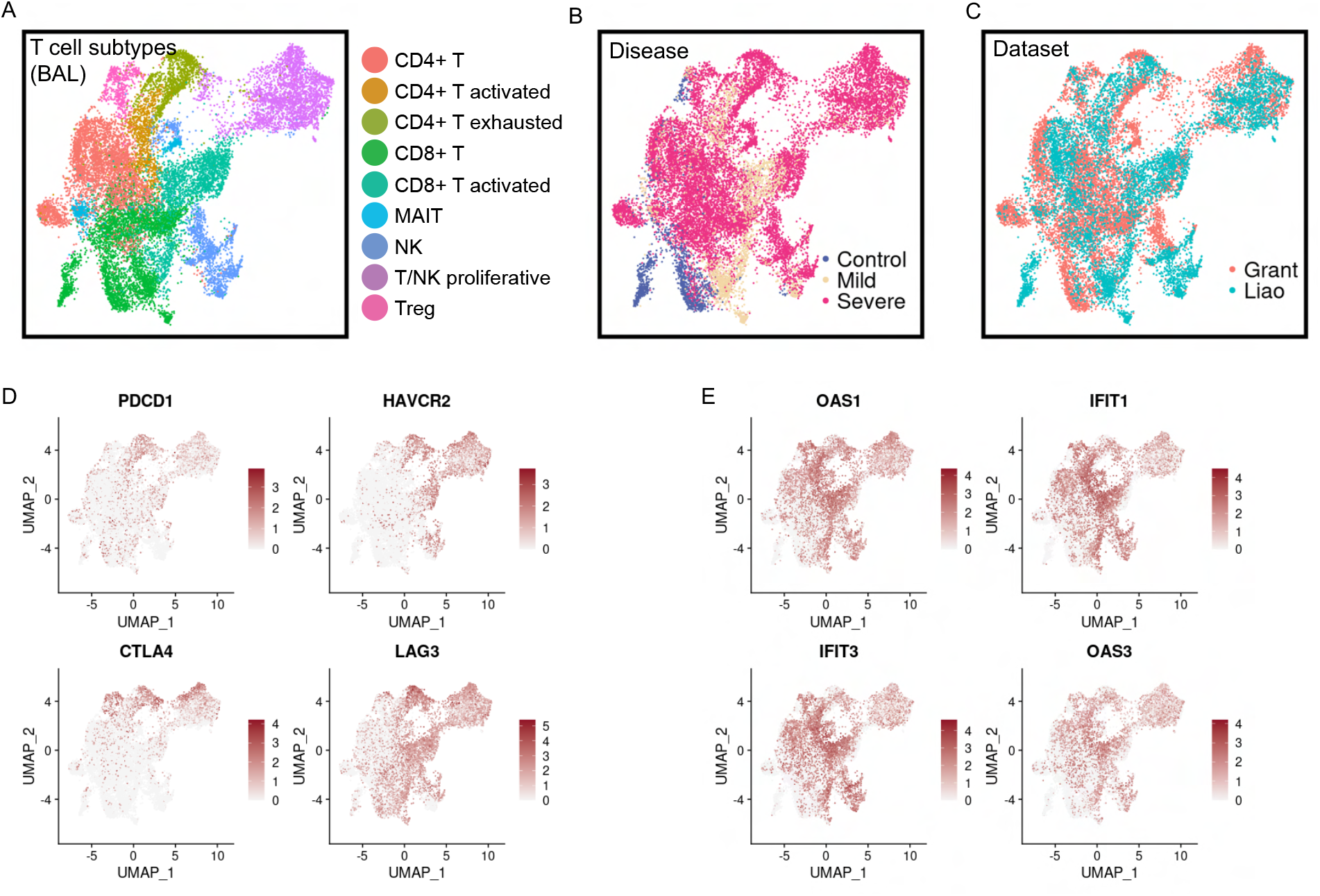
Distinct subtypes of T cells and NK cells in COVID-19 BAL data. (**A-C**) UMAPs of subtypes (A), COVID-19 conditions (B) and data sources (C) of T cells and NK cells in the integrated BAL data. (D-E) UMAPs of normalized expression values of exhausted T cell markers (**D**) and ISGs (**E**).

**Figure S17.**
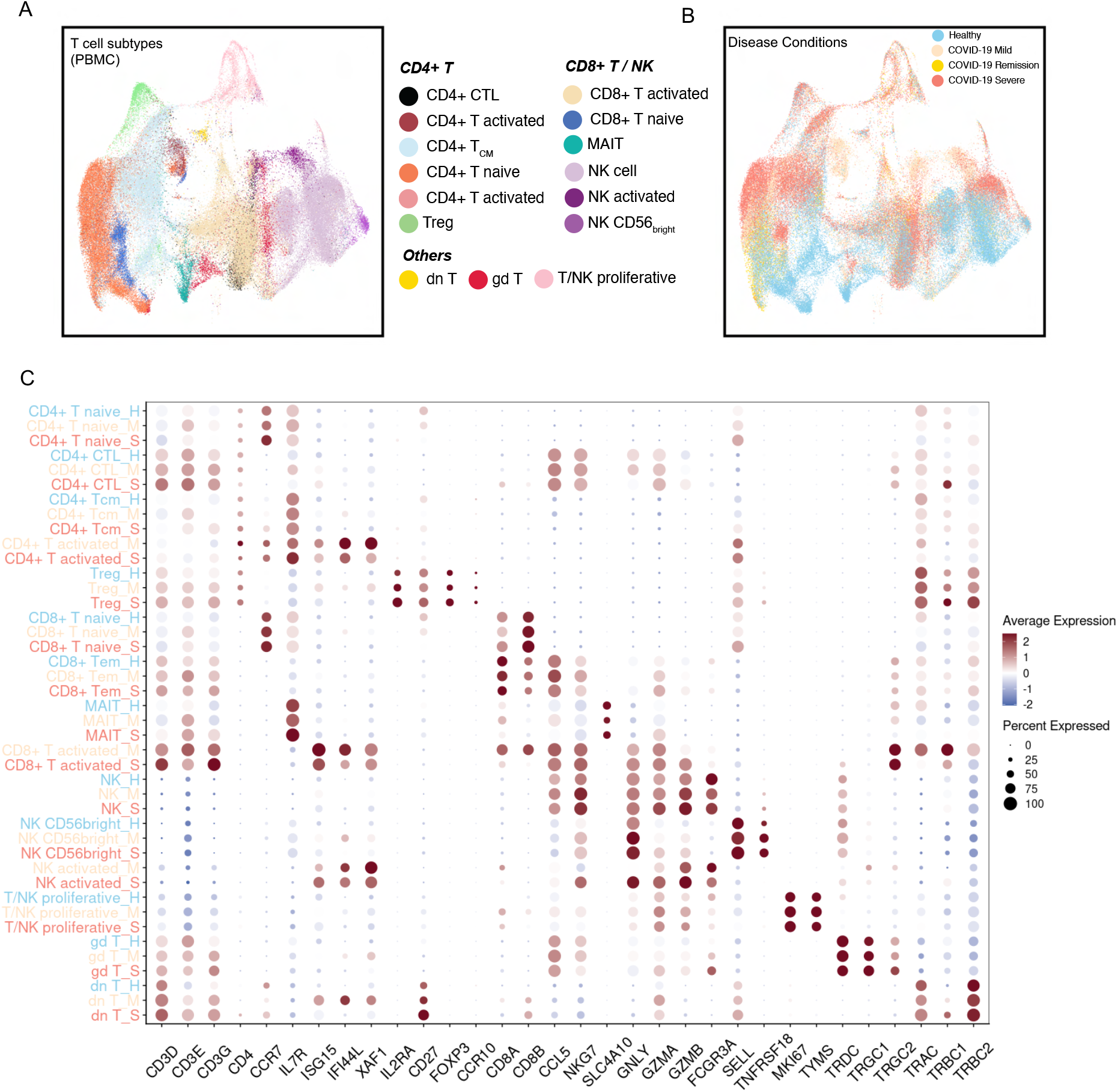
Various T cell and NK cell subtypes in the integrated PBMC data. (**A-B**) UMAPs of T cell and NK cell subtypes (A) and COVID-19 conditions (B) after integration of T cells in 5 PBMC single-cell datasets. (**C**) Dot plot shows T cell and NK cell subtype associated genes for each subtype per disease condition. Labels of cell types of healthy donors, mild patients and severe patients are colored by blue, yellow and red. Scaled expression values are shown using a color scheme.

**Figure S18.**
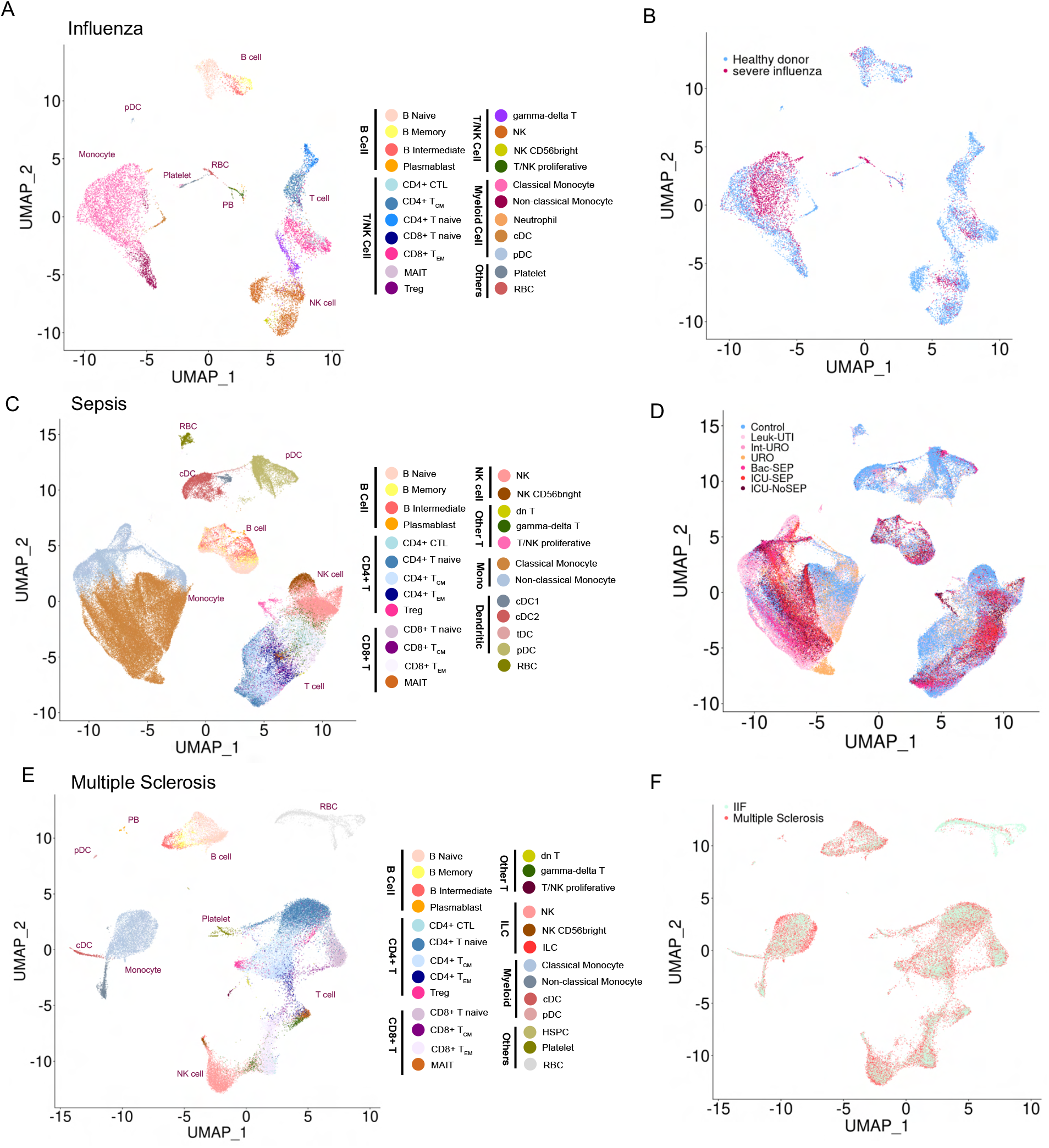
Various cell types in immune-mediated diseases, relative to Figure 7. (**A, C, E**) Distributions of cell types identified in influenza (A), sepsis (C) and multiple sclerosis (E) patients were shown on UMAPs. (**B, D, F**) Distributions of disease conditions in influenza (B), sepsis (D) and multiple sclerosis (F) patients were shown on UMAPs.

## Notes

https://toppcell.cchmc.org/biosystems/go/index3/COVID-19%20Atlas

### Competing Interest Statement

The authors have declared no competing interest.

### Summary of Updates

This version improves the figure descriptions (legends) and also distinguishes more succinctly the description of the web resource itself versus how users can develop hypotheses through the use of the system versus the specifics of the three principle hypotheses that we develop as examples of using the system.

## Reference

Aid, M., Busman-Sahay, K., Vidal, S.J., Maliga, Z., Bondoc, S., Starke, C., Terry, M., Jacobson, C.A., Wrijil, L., Ducat, S., et al. (2020). Vascular Disease and Thrombosis in SARS-CoV-2-Infected Rhesus Macaques. Cell 183, 1354–1366.e13.

Arazi, A., Rao, D.A., Berthier, C.C., Davidson, A., Liu, Y., Hoover, P.J., Chicoine, A., Eisenhaure, T.M., Jonsson, A.H., Li, S., et al. (2019). The immune cell landscape in kidneys of patients with lupus nephritis. Nat. Immunol. 20, 902–914.

Arunachalam, P.S., Wimmers, F., Mok, C.K.P., Perera, R.A.P.M., Scott, M., Hagan, T., Sigal, N., Feng, Y., Bristow, L., Tak-Yin Tsang, O., et al. (2020). Systems biological assessment of immunity to mild versus severe COVID-19 infection in humans. Science 369, 1210–1220.

Aust, G., Sittig, D., Becherer, L., Anderegg, U., Schütz, A., Lamesch, P., and Schmücking, E. (2004). The role of CXCR5 and its ligand CXCL13 in the compartmentalization of lymphocytes in thyroids affected by autoimmune thyroid diseases. Eur. J. Endocrinol. 150, 225–234.

Barnes, B.J., Adrover, J.M., Baxter-Stoltzfus, A., Borczuk, A., Cools-Lartigue, J., Crawford, J.M., Daßler-Plenker, J., Guerci, P., Huynh, C., Knight, J.S., et al. (2020). Targeting potential drivers of COVID-19: Neutrophil extracellular traps. J. Exp. Med. 217.

Bharat, A., Querrey, M., Markov, N.S., Kim, S., Kurihara, C., Garza-Castillon, R., Manerikar, A., Shilatifard, A., Tomic, R., Politanska, Y., et al. (2020). Lung transplantation for patients with severe COVID-19. Sci. Transl. Med. 12.

Blanco-Melo, D., Nilsson-Payant, B.E., Liu, W.-C., Uhl, S., Hoagland, D., Møller, R., Jordan, T.X., Oishi, K., Panis, M., Sachs, D., et al. (2020). Imbalanced Host Response to SARS-CoV-2 Drives Development of COVID-19. Cell 181, 1036–1045.e9.

Blighe, K., Rana, S., and Lewis, M. (2018). EnhancedVolcano: Publication-Ready Volcano Plots With Enhanced Colouring and Labeling.(2019). R Package Version 1.

Cao, X. (2020). COVID-19: immunopathology and its implications for therapy. Nat. Rev. Immunol. 20, 269–270.

Chua, R.L., Lukassen, S., Trump, S., Hennig, B.P., Wendisch, D., Pott, F., Debnath, O., Thürmann, L., Kurth, F., Völker, M.T., et al. (2020). COVID-19 severity correlates with airway epithelium–immune cell interactions identified by single-cell analysis. Nat. Biotechnol. 38, 970–979.

De Biasi, S., Lo Tartaro, D., Meschiari, M., Gibellini, L., Bellinazzi, C., Borella, R., Fidanza, L., Mattioli, M., Paolini, A., Gozzi, L., et al. (2020). Expansion of plasmablasts and loss of memory B cells in peripheral blood from COVID-19 patients with pneumonia. Eur. J. Immunol. 50, 1283–1294.

Delorey, T.M., Ziegler, C.G.K., Heimberg, G., Normand, R., Yang, Y., Segerstolpe, A., Abbondanza, D., Fleming, S.J., Subramanian, A., Montoro, D.T., et al. (2021). A single-cell and spatial atlas of autopsy tissues reveals pathology and cellular targets of SARS-CoV-2. bioRxiv.

Edovitsky, E., Lerner, I., Zcharia, E., Peretz, T., Vlodavsky, I., and Elkin, M. (2006). Role of endothelial heparanase in delayed-type hypersensitivity. Blood 107, 3609–3616.

Ehrenfeld, M., Tincani, A., Andreoli, L., Cattalini, M., Greenbaum, A., Kanduc, D., Alijotas-Reig, J., Zinserling, V., Semenova, N., Amital, H., et al. (2020). Covid-19 and autoimmunity. Autoimmun. Rev. 19, 102597.

Farris, A.D., and Guthridge, J.M. (2020). Overlapping B cell pathways in severe COVID-19 and lupus. Nat. Immunol. 21, 1478–1480.

Fujii, H., Tsuji, T., Yuba, T., Tanaka, S., Suga, Y., Matsuyama, A., Omura, A., Shiotsu, S., Takumi, C., Ono, S., et al. (2020). High levels of anti-SSA/Ro antibodies in COVID-19 patients with severe respiratory failure: a case-based review. Clin. Rheumatol. 39, 3171–3175.

Goodall, K.J., Poon, I.K.H., Phipps, S., and Hulett, M.D. (2014). Soluble heparan sulfate fragments generated by heparanase trigger the release of pro-inflammatory cytokines through TLR-4. PLoS One 9, e109596.

Grant, R.A., Morales-Nebreda, L., and Markov, N.S. (2020). Alveolitis in severe SARS-CoV-2 pneumonia is driven by self-sustaining circuits between infected alveolar macrophages and T cells. bioRxiv.

Gruber, C.N., Patel, R.S., Trachtman, R., Lepow, L., Amanat, F., Krammer, F., Wilson, K.M., Onel, K., Geanon, D., Tuballes, K., et al. (2020). Mapping Systemic Inflammation and Antibody Responses in Multisystem Inflammatory Syndrome in Children (MIS-C). Cell 183, 982–995.e14.

Guo, C., Li, B., Ma, H., Wang, X., Cai, P., Yu, Q., Zhu, L., Jin, L., Jiang, C., Fang, J., et al. (2020). Single-cell analysis of two severe COVID-19 patients reveals a monocyte-associated and tocilizumab-responding cytokine storm. Nat. Commun. 11, 3924.

Hadjadj, J., Yatim, N., Barnabei, L., Corneau, A., Boussier, J., Smith, N., Péré, H., Charbit, B., Bondet, V., Chenevier-Gobeaux, C., et al. (2020). Impaired type I interferon activity and inflammatory responses in severe COVID-19 patients. Science 369, 718–724.

Hao, Y., Hao, S., Andersen-Nissen, E., Mauck, W.M., Zheng, S., Butler, A., Lee, M.J., Wilk, A.J., Darby, C., Zagar, M., et al. (2020). Integrated analysis of multimodal single-cell data.

He, R., Lu, Z., Zhang, L., Fan, T., Xiong, R., Shen, X., Feng, H., Meng, H., Lin, W., Jiang, W., et al. (2020). The clinical course and its correlated immune status in COVID-19 pneumonia. J. Clin. Virol. 127, 104361.

Heemskerk, J.W.M., Mattheij, N.J.A., and J M E (2013). Platelet-based coagulation: different populations, different functions. Journal of Thrombosis and Haemostasis 11, 2–16.

Heming, M., Li, X., Räuber, S., Mausberg, A.K., Börsch, A.-L., Hartlehnert, M., Singhal, A., Lu, I.-N., Fleischer, M., Szepanowski, F., et al. (2021). Neurological Manifestations of COVID-19 Feature T Cell Exhaustion and Dedifferentiated Monocytes in Cerebrospinal Fluid. Immunity 54, 164–175.e6.

Hirota, K., Yoshitomi, H., Hashimoto, M., Maeda, S., Teradaira, S., Sugimoto, N., Yamaguchi, T., Nomura, T., Ito, H., Nakamura, T., et al. (2007). Preferential recruitment of CCR6-expressing Th17 cells to inflamed joints via CCL20 in rheumatoid arthritis and its animal model. J. Exp. Med. 204, 2803–2812.

Iba, T., Levy, J.H., Connors, J.M., Warkentin, T.E., Thachil, J., and Levi, M. (2020a). The unique characteristics of COVID-19 coagulopathy. Crit. Care 24, 360.

Iba, T., Connors, J.M., and Levy, J.H. (2020b). The coagulopathy, endotheliopathy, and vasculitis of COVID-19. Inflamm. Res. 69, 1181–1189.

Jayatilleke, K.M., and Hulett, M.D. (2020). Heparanase and the hallmarks of cancer. J. Transl. Med. 18, 453.

Jin, S., Guerrero-Juarez, C.F., Zhang, L., Chang, I., Myung, P., Plikus, M.V., and Nie, Q. (2020). Inference and analysis of cell-cell communication using CellChat.

Kaimal, V., Bardes, E.E., Tabar, S.C., Jegga, A.G., and Aronow, B.J. (2010). ToppCluster: a multiple gene list feature analyzer for comparative enrichment clustering and network-based dissection of biological systems. Nucleic Acids Res. 38, W96–W102.

Klimatcheva, E., Pandina, T., Reilly, C., Torno, S., Bussler, H., Scrivens, M., Jonason, A., Mallow, C., Doherty, M., Paris, M., et al. (2015). CXCL13 antibody for the treatment of autoimmune disorders. BMC Immunol. 16, 6.

Kuwabara, T., Ishikawa, F., Yasuda, T., Aritomi, K., Nakano, H., Tanaka, Y., Okada, Y., Lipp, M., and Kakiuchi, T. (2009). CCR 7 Ligands Are Required for Development of Experimental Autoimmune Encephalomyelitis through Generating IL-23-Dependent Th17 Cells. The Journal of Immunology 183, 2513–2521.

Laing, A.G., Lorenc, A., Del Molino Del Barrio, I., Das, A., Fish, M., Monin, L., Muñoz-Ruiz, M., McKenzie, D.R., Hayday, T.S., Francos-Quijorna, I., et al. (2020). Author Correction: A dynamic COVID-19 immune signature includes associations with poor prognosis. Nat. Med. 26, 1951.

Lee, H.-T., Shiao, Y.-M., Wu, T.-H., Chen, W.-S., Hsu, Y.-H., Tsai, S.-F., and Tsai, C.-Y. (2010). Serum BLC/CXCL13 concentrations and renal expression of CXCL13/CXCR5 in patients with systemic lupus erythematosus and lupus nephritis. J. Rheumatol. 37, 45–52.

Lee, J.S., Park, S., Jeong, H.W., Ahn, J.Y., Choi, S.J., Lee, H., Choi, B., Nam, S.K., Sa, M., Kwon, J.-S., et al. (2020). Immunophenotyping of COVID-19 and influenza highlights the role of type I interferons in development of severe COVID-19. Sci Immunol 5.

Levi, M., Thachil, J., Iba, T., and Levy, J.H. (2020). Coagulation abnormalities and thrombosis in patients with COVID-19. The Lancet Haematology 7, e438–e440.

Liao, M., Liu, Y., Yuan, J., Wen, Y., Xu, G., Zhao, J., Cheng, L., Li, J., Wang, X., Wang, F., et al. (2020). Single-cell landscape of bronchoalveolar immune cells in patients with COVID-19. Nat. Med. 26, 842–844.

Mason, D.Y., Cordell, J.L., Brown, M.H., Borst, J., Jones, M., Pulford, K., Jaffe, E., Ralfkiaer, E., Dallenbach, F., and Stein, H. (1995). CD79a: a novel marker for B-cell neoplasms in routinely processed tissue samples. Blood 86, 1453–1459.

McGonagle, D., Sharif, K., O’Regan, A., and Bridgewood, C. (2020). The Role of Cytokines including Interleukin-6 in COVID-19 induced Pneumonia and Macrophage Activation Syndrome-Like Disease. Autoimmun. Rev. 19, 102537.

Mehta, P., McAuley, D.F., Brown, M., Sanchez, E., Tattersall, R.S., Manson, J.J., and HLH Across Speciality Collaboration, UK (2020). COVID-19: consider cytokine storm syndromes and immunosuppression. Lancet 395, 1033–1034.

Merad, M., and Martin, J.C. (2020). Author Correction: Pathological inflammation in patients with COVID-19: a key role for monocytes and macrophages. Nat. Rev. Immunol. 20, 448.

Middleton, E.A., He, X.-Y., Denorme, F., Campbell, R.A., Ng, D., Salvatore, S.P., Mostyka, M., Baxter-Stoltzfus, A., Borczuk, A.C., Loda, M., et al. (2020). Neutrophil extracellular traps contribute to immunothrombosis in COVID-19 acute respiratory distress syndrome. Blood 136, 1169–1179.

Nicolai, L., Leunig, A., Brambs, S., Kaiser, R., Weinberger, T., Weigand, M., Muenchhoff, M., Hellmuth, J.C., Ledderose, S., Schulz, H., et al. (2020). Immunothrombotic Dysregulation in COVID-19 Pneumonia Is Associated With Respiratory Failure and Coagulopathy. Circulation 142, 1176–1189.

Osterholm, C., Folkersen, L., Lengquist, M., Pontén, F., Renné, T., Li, J., and Hedin, U. (2013). Increased expression of heparanase in symptomatic carotid atherosclerosis. Atherosclerosis 226, 67–73.

Pedersen, S.F., and Ho, Y.-C. (2020). SARS-CoV-2: a storm is raging. J. Clin. Invest. 130, 2202–2205.

Rapkiewicz, A.V., Mai, X., Carsons, S.E., Pittaluga, S., Kleiner, D.E., Berger, J.S., Thomas, S., Adler, N.M., Charytan, D.M., Gasmi, B., et al. (2020). Megakaryocytes and platelet-fibrin thrombi characterize multi-organ thrombosis at autopsy in COVID-19: A case series. EClinicalMedicine 24, 100434.

Reyes, M., Filbin, M.R., Bhattacharyya, R.P., Billman, K., Eisenhaure, T., Hung, D.T., Levy, B.D., Baron, R.M., Blainey, P.C., Goldberg, M.B., et al. (2020). An immune-cell signature of bacterial sepsis. Nat. Med. 26, 333–340.

Rodríguez, Y., Novelli, L., Rojas, M., De Santis, M., Acosta-Ampudia, Y., Monsalve, D.M., Ramírez-Santana, C., Costanzo, A., Ridgway, W.M., Ansari, A.A., et al. (2020). Autoinflammatory and autoimmune conditions at the crossroad of COVID-19. J. Autoimmun. 114, 102506.

Schafflick, D., Xu, C.A., Hartlehnert, M., Cole, M., Schulte-Mecklenbeck, A., Lautwein, T., Wolbert, J., Heming, M., Meuth, S.G., Kuhlmann, T., et al. (2020). Integrated single cell analysis of blood and cerebrospinal fluid leukocytes in multiple sclerosis. Nature Communications 11.

Schulte-Schrepping, J., Reusch, N., Paclik, D., Baßler, K., Schlickeiser, S., Zhang, B., Krämer, B., Krammer, T., Brumhard, S., Bonaguro, L., et al. (2020). Severe COVID-19 Is Marked by a Dysregulated Myeloid Cell Compartment. Cell 182, 1419–1440.e23.

Schultz, N.H., Sørvoll, I.H., Michelsen, A.E., Munthe, L.A., Lund-Johansen, F., Ahlen, M.T., Wiedmann, M., Aamodt, A.-H., Skattør, T.H., Tjønnfjord, G.E., et al. (2021). Thrombosis and Thrombocytopenia after ChAdOx1 nCoV-19 Vaccination. N. Engl. J. Med.

Shi, Y., Wang, Y., Shao, C., Huang, J., Gan, J., Huang, X., Bucci, E., Piacentini, M., Ippolito, G., and Melino, G. (2020). COVID-19 infection: the perspectives on immune responses. Cell Death Differ. 27, 1451–1454.

Silvin, A., Chapuis, N., Dunsmore, G., Goubet, A.-G., Dubuisson, A., Derosa, L., Almire, C., Hénon, C., Kosmider, O., Droin, N., et al. (2020). Elevated Calprotectin and Abnormal Myeloid Cell Subsets Discriminate Severe from Mild COVID-19. Cell 182, 1401–1418.e18.

Sparkenbaugh, E., and Pawlinski, R. (2013). Interplay between coagulation and vascular inflammation in sickle cell disease. Br. J. Haematol. 162, 3–14.

Spoerl, S., Kremer, A.N., Aigner, M., Eisenhauer, N., Koch, P., Meretuk, L., Löffler, P., Tenbusch, M., Maier, C., Überla, K., et al. (2021). Upregulation of CCR4 in activated CD8+ T cells indicates enhanced lung homing in patients with severe acute SARS-CoV-2 infection. Eur. J. Immunol.

Steinmetz, O.M., Velden, J., Kneissler, U., Marx, M., Klein, A., Helmchen, U., Stahl, R.A.K., and Panzer, U. (2008). Analysis and classification of B-cell infiltrates in lupus and ANCA-associated nephritis. Kidney Int. 74, 448–457.

Stuart, T., Butler, A., Hoffman, P., Hafemeister, C., Papalexi, E., Mauck, W.M., 3rd, Hao, Y., Stoeckius, M., Smibert, P., and Satija, R. (2019). Comprehensive Integration of Single-Cell Data. Cell 177, 1888–1902.e21.

Swieringa, F., Spronk, H.M.H., Heemskerk, J.W.M., and van der Meijden, P.E.J. (2018). Integrating platelet and coagulation activation in fibrin clot formation. Res Pract Thromb Haemost 2, 450–460.

Tay, M.Z., Poh, C.M., Rénia, L., MacAry, P.A., and Ng, L.F.P. (2020). The trinity of COVID-19: immunity, inflammation and intervention. Nat. Rev. Immunol. 20, 363–374.

Terpos, E., Ntanasis-Stathopoulos, I., Elalamy, I., Kastritis, E., Sergentanis, T.N., Politou, M., Psaltopoulou, T., Gerotziafas, G., and Dimopoulos, M.A. (2020). Hematological findings and complications of COVID-19. Am. J. Hematol. 95, 834–847.

Thålin, C., Hisada, Y., Lundström, S., Mackman, N., and Wallén, H. (2019). Neutrophil Extracellular Traps: Villains and Targets in Arterial, Venous, and Cancer-Associated Thrombosis. Arterioscler. Thromb. Vasc. Biol. 39, 1724–1738.

Wang, F., Nie, J., Wang, H., Zhao, Q., Xiong, Y., Deng, L., Song, S., Ma, Z., Mo, P., and Zhang, Y. (2020). Characteristics of Peripheral Lymphocyte Subset Alteration in COVID-19 Pneumonia. J. Infect. Dis. 221, 1762–1769.

Wilk, A.J., Rustagi, A., Zhao, N.Q., Roque, J., Martínez-Colón, G.J., McKechnie, J.L., Ivison, G.T., Ranganath, T., Vergara, R., Hollis, T., et al. (2020a). A single-cell atlas of the peripheral immune response in patients with severe COVID-19. Nature Medicine 26, 1070–1076.

Wilk, A.J., Lee, M.J., Wei, B., Parks, B., Pi, R., Martínez-Colón, G.J., Ranganath, T., Zhao, N.Q., Taylor, S., Becker, W., et al. (2020b). Multi-omic profiling reveals widespread dysregulation of innate immunity and hematopoiesis in COVID-19.

Wolf, F.A., Angerer, P., and Theis, F.J. (2018). SCANPY: large-scale single-cell gene expression data analysis. Genome Biol. 19, 15.

Wong, C.K., Wong, P.T.Y., Tam, L.S., Li, E.K., Chen, D.P., and Lam, C.W.K. (2010). Elevated production of B cell chemokine CXCL13 is correlated with systemic lupus erythematosus disease activity. J. Clin. Immunol. 30, 45–52.

Yang, A.C., Kern, F., Losada, P.M., Maat, C.A., and Schmartz, G. (2020). Broad transcriptional dysregulation of brain and choroid plexus cell types with COVID-19. bioRxiv.

Zhang, F., Wei, K., Slowikowski, K., Fonseka, C.Y., Rao, D.A., Kelly, S., Goodman, S.M., Tabechian, D., Hughes, L.B., Salomon-Escoto, K., et al. (2019). Defining inflammatory cell states in rheumatoid arthritis joint synovial tissues by integrating single-cell transcriptomics and mass cytometry. Nat. Immunol. 20, 928–942.

Zhao, M. (2020). Cytokine storm and immunomodulatory therapy in COVID-19: Role of chloroquine and anti-IL-6 monoclonal antibodies. Int. J. Antimicrob. Agents 55, 105982.

Zhou, Y., Han, T., Chen, J., Hou, C., Hua, L., He, S., Guo, Y., Zhang, S., Wang, Y., Yuan, J., et al. (2020). Clinical and autoimmune characteristics of severe and critical cases of COVID-19. Clin. Transl. Sci. 13, 1077–1086.

Zuo, Y., Zuo, M., Yalavarthi, S., Gockman, K., Madison, J.A., Shi, H., Woodard, W., Lezak, S.P., Lugogo, N.L., Knight, J.S., et al. (2021). Neutrophil extracellular traps and thrombosis in COVID-19. J. Thromb. Thrombolysis 51, 446–453.

